# L-Type Ca^2+^ channels and TRPC3 channels shape brain pericyte Ca^2+^ signaling and hemodynamics throughout the arteriole to capillary network *in vivo*

**DOI:** 10.1101/2024.02.27.582351

**Authors:** Jessica Meza-Resillas, Finnegan O’Hara, Syed Kaushik, Michael Stobart, Noushin Ahmadpour, Meher Kantroo, Shahin Shabanipour, John Del Rosario, Megan C. Rodriguez, Dmytro Koval, Chaim Glück, Bruno Weber, Jillian Stobart

## Abstract

Pericytes play a crucial role in regulating cerebral blood flow (CBF) through processes like vasomotion and neurovascular coupling (NVC). Recent work has identified different pericyte types at distinct points in the cerebrovascular network, such as the arteriole-capillary transition zone (ACT) and distal capillaries, sparking debate about their functional roles in blood flow control. Part of this discussion has comprised the possible mechanisms that may regulate pericyte Ca^2+^ signaling. Using *in vivo* two-photon Ca^2+^ imaging and a pharmacological approach with Ca^2+^ channel blockers (nimodipine and Pyr3), we assessed the contribution of L-type voltage-gated Ca^2+^ channels (VGCC) and transient receptor potential canonical 3 (TRPC3) channels to Ca^2+^ signaling in different pericyte types, ensheathing and capillary pericytes. We also measured local hemodynamics such as vessel diameter, blood cell velocity and flux during vasomotion, and following somatosensory stimulation to evoke NVC. We report that VGCC and TRPC3 channels underlie spontaneous fluctuations in ensheathing pericyte Ca^2+^ that trigger vasomotor contractions, but the contribution of each of these mechanisms to vascular tone depends on the specific branch of the ACT. Distal capillary pericytes also express L-type VGCCs and TRPC3 channels and they mediate spontaneous Ca^2+^ signaling in these cells. However, only TRPC3 channels maintain resting capillary tone, possibly by a receptor-operated Ca^2+^ entry mechanism. By applying the Ca^2+^ channel blockers during NVC, we found a significant involvement of L-type VGCCs in both pericyte types, influencing their ability to dilate during functional hyperemia. These findings provide new evidence of VGCC and TRPC3 activity in pericytes *in vivo* and establish a clear distinction between brain pericyte types and their functional roles, opening avenues for innovative strategies to selectively target their Ca^2+^ dynamics for CBF control.

**Significance Statement:** Although brain pericytes contribute to the regulation of CBF, there is uncertainty about how different types of pericytes are involved in this process. Ca^2+^ signaling is believed to be important for the contractility and tone of pericytes, but there is a limited understanding of the Ca^2+^ pathways in specific pericyte types. Here, we demonstrate that both VGCC and TRPC3 channels are active in distinct types of pericytes throughout the cerebrovascular network, but have different roles in pericyte tone depending on the pericyte location. This has important implications for how pericytes influence vasomotion and neurovascular coupling, which are central processes in CBF regulation. This work also provides the first evidence of TRPC3 channel activity in pericytes *in vivo*, furthering our understanding of the diverse signaling pathways within these brain mural cells.

## Introduction

The neurovascular unit is comprised of neurons, glia, smooth muscle cells, endothelial cells and pericytes (1, 2). These cells collaborate to ensure proper blood delivery to the brain during processes like vasomotion and neurovascular coupling. Vasomotion refers to spontaneous, rhythmic contraction and dilation events in blood vessels. As a result, this phenomenon induces blood flow motion, facilitating its delivery within an organ (3). Neurovascular coupling is the dynamic temporal and regional adjustment of CBF in response to neuronal activity and metabolic signals (4), which is crucial for adequate supply of blood to support information processing and neuronal activity (5, 6). NVC and vasomotion represent pivotal physiological processes in which pericytes, assume a paramount role (7–12).

Brain pericytes are situated within the basolateral membrane of brain blood vessels and have a protruding soma (bump-on-a-log shape) and processes that extend along the vessels (8, 13). Pericytes at different points within the brain vascular network can be sub-classified based on their morphology, protein expression, and potential functional roles (13, 14). Ensheathing pericytes are located in the arteriole-capillary transition zone (ACT) from the 1^st^ to 4^th^ branches after the penetrating arteriole (14). These cells express alpha-smooth muscle actin (α-SMA) and have ovoid somata with processes that enwrap blood vessels (13, 14). They contribute significantly to the regulation of CBF (7, 15–22), and dilate quickly during NVC to supply blood downstream (23). Deeper in the vascular network, distal capillary pericytes are found 5 or more branches from the penetrating arteriole and extend their processes along capillaries (13, 14). These pericytes have little to no expression of α-SMA, but they have the capacity to constrict capillaries (24). This suggests they maintain capillary tone and resistance, and may play a role in NVC.

Both ensheathing and capillary pericytes express a broad range of Ca^2+^ channels and vasomodulator receptors that can trigger fluctuations in intracellular Ca^2+^. Elevated intracellular Ca^2+^ in ensheathing pericytes causes cellular contraction (25, 26), and this is highly mediated by L-type VGCC activity (7, 15, 22, 26). In distal capillary pericytes, the activity of VGCC appears to be less robust than in ensheathing pericytes, since VGCC blockers mildly reduce or do not affect spontaneous Ca^2+^ signaling in capillary pericytes in brain slices (25, 26), and these channels are not activated during pressure-induced capillary constriction unlike upstream ensheathing pericytes and smooth muscle cells (22). However, VGCC are reportedly expressed by capillary pericytes (27, 28), and the contribution of these channels to capillary pericyte Ca^2+^ signaling and cortical hemodynamics *in vivo* remains unknown. Other Ca^2+^ channels also contribute to capillary pericyte Ca^2+^ events, such as store operated Ca^2+^ entry channels (26) and transient receptor potential (TRP) channels (25, 26). Specifically, TRPC3 channels are expressed in brain pericytes as determined by transcriptomics (27), but their protein expression, contribution to Ca^2+^ signaling in the pericyte subtypes, and influence on blood vessel hemodynamics have not been confirmed. TRPC3 channels can amplify intracellular Ca^2+^ triggered by Gq-GPCR signaling as a receptor-operated Ca^2+^ entry pathway (29–31), which is potentially important for pericyte Ca^2+^ events.

Here, we used pharmacological tools to investigate the contribution of L-type VGCC and TRPC3 channels to Ca^2+^ signaling in ensheathing and capillary pericytes *in vivo* and the impact of this signaling on local hemodynamics during vasomotion and NVC. Our results suggest that each of these Ca^2+^ channels have distinct roles in ensheathing and capillary pericytes that affects blood flow control at specific points of the brain vasculature.

## Results

### Brain pericytes express L-type VGCC and TRPC3 channels

We classified different mural cell populations based on their location within the vascular network and morphology as described above. Ensheathing pericytes were located in the ACT (1^st^ to 4^th^ branches after the penetrating arteriole) and covered blood vessels with their processes (Fig. 1A)(14). Capillary pericytes were found 5 or more branches from the penetrating arteriole and displayed a “thin-strand” morphology (13, 14). We identified both pericyte types in fixed cortical sections by CD13 (mural cell) and CD31 (endothelial cell) immunohistochemistry (Fig. 1). Antibodies for the Cav 1.2 subunit of L-type channels and TRPC3 channels confirmed the expression of these proteins in both ensheathing and capillary pericytes (Fig. 1B-D).

**Figure 1.**
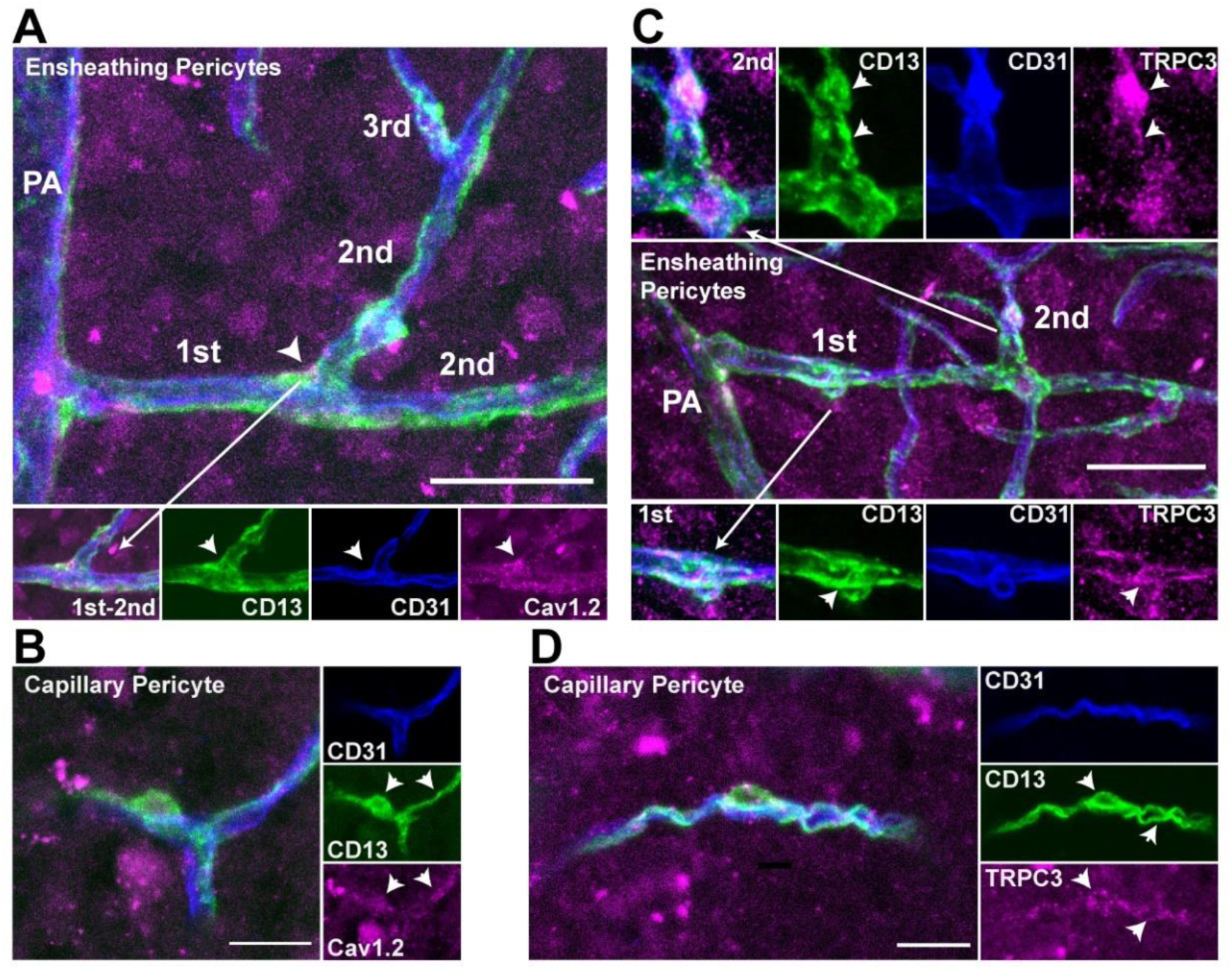
Expression of L-type VGCC and TRPC3 channels by cortical ensheathing and capillary pericytes. **A)** Immunohistochemistry shows the antibody for Cav1.2, a subunit of L-type VGCC, localized to ensheathing pericytes (anti-CD13) near endothelial cells (anti-CD31), in branches of the ACT. Arrowhead indicates the pericyte soma. Scale = 25 µm. **B)** Cav1.2 staining also localizes to capillary pericytes found more than 5 branches from the penetrating arteriole. Arrowheads indicate the soma and process. Scale = 10 µm. **C)** The antibody for TRPC3 channels also localizes to ensheathing pericytes in the ACT. Arrowheads indicate the somata and processes. Scale = 25 µm. D) TRPC3 staining also localizes to capillary pericytes (anti-CD13). Arrowheads indicate soma and process. Scale = 10 µm.

### L-type VGCC and TRPC3 channel blockers affect ensheathing pericyte Ca^2+^ and resting blood vessel hemodynamics in the arteriole-capillary transition zone

To study the role of VGCC and TRPC3 channels in ensheathing pericytes *in vivo*, we utilized *Acta2*-RCaMP1.07 mice (Fig. 2A). Due to the α-SMA promoter, the red genetically encoded Ca^2+^ indicator RCaMP1.07 was expressed in vascular smooth muscle cells on penetrating arterioles and ensheathing pericytes in the ACT (Fig. 2B), as we observed by two-photon microscopy through a chronic cranial window in ketamine/xylazine anesthetized mice. We recorded basal fluctuations in ensheathing pericytes Ca^2+^ at baseline and following i.p. injection of nimodipine (L-type VGCC blocker; 1 mg/kg) or Pyr3 (TRPC3 channel blocker; 20 mg/kg) which reportedly cross the blood-brain-barrier (32–34). Data was acquired from the same pericytes in different imaging sessions for each drug. We found that the amplitude and frequency of Ca^2+^ signals in ensheathing pericyte somata and processes were reduced by nimodipine (Fig. 2C-E). Pyr3 decreased the frequency of Ca^2+^ events in ensheathing pericyte processes but did not affect the amplitude (Fig. 2C-E).

**Figure 2.**
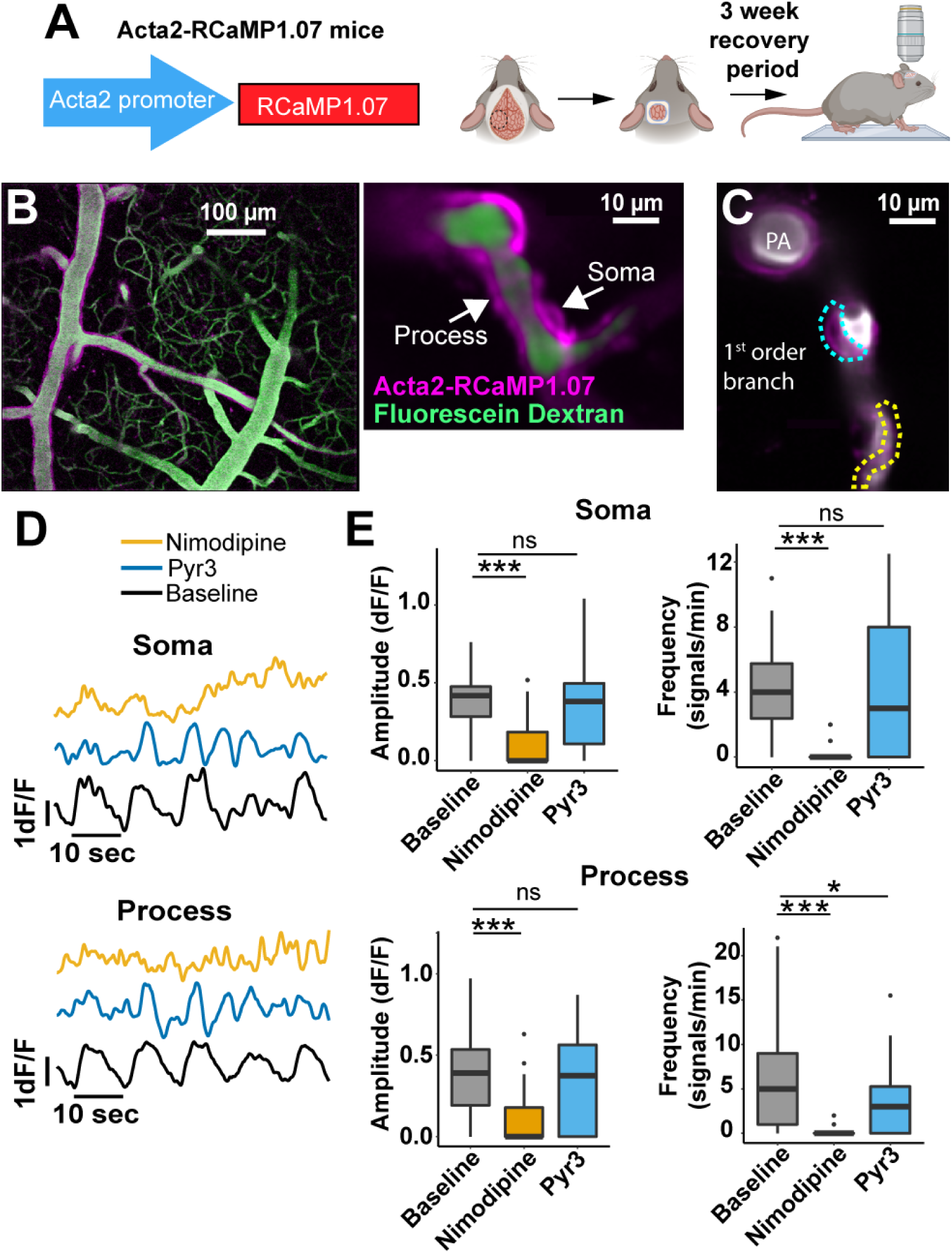
Ensheathing pericyte Ca^2+^ imaging during treatment with L-type VGCC and TRPC3 channel blockers. **A)** *Acta2*-RCaMP1.07 mouse model and chronic cranial window surgical scheme. **B)** Left: Visualization of ACT mural cells expressing RCaMP1.07 (magenta) under αSMA promoter labeled ensheathing pericytes (right). The vasculature was labeled with fluorescein dextran (i.v.; green). **C)** Two-photon image of ensheathing pericyte morphological structures-Soma: cyan dashed line; Process; yellow dashed line. **D)** Individual Ca^2+^ signaling traces of soma and process from C). **E)** Spontaneous Ca^2+^ signaling properties (amplitude and frequency) from ensheathing pericyte somata and processes. n=43 pericytes from 7 mice. For specific p-values and mean ± SD information, please refer to SI Table S1.1 and S1.2.

In the same animals where Ca^2+^ was measured, we labeled the blood with fluorescein dextran (2.5 %, i.v.) and conducted two-photon line scans to measure blood vessel diameter, blood cell (BC) velocity and flux. Again, the same vessels were measured across multiple imaging sessions at baseline and following nimodipine or Pyr3 injection. Both nimodipine and Pyr3 caused a dilatory effect in blood vessels covered by ensheathing pericytes (Fig. 3A-C). Considering that the blood vessels of the ACT have vasomotion (14, 25), we also examined the diameter amplitude change (ΔD/D) which we termed the vasomotor index, as well as the vasomotor frequency that gives the number of vessel oscillations per minute. Nimodipine decreased both characteristics of vasomotion (Fig. 3D, E), while Pyr3 reduced the vasomotor index, but did not alter the frequency of vasomotor oscillations (Fig. 3D, E). Next, we assessed the impact of the Ca^2+^ channel blockers on BC dynamics. Nimodipine decreased BC velocity (Fig. 3H) and occasionally caused brief stalling of blood flow (Fig. 3F, G). This did not alter the BC flux (Fig. 3I, J). Conversely, Pyr3 tended to increase velocity (p-value=0.067), and significantly increased BC flux (Fig. 3H, J). This suggests that both VGCC and TRPC3 contribute to ensheathing pericyte contractility and local hemodynamics.

**Figure 3.**
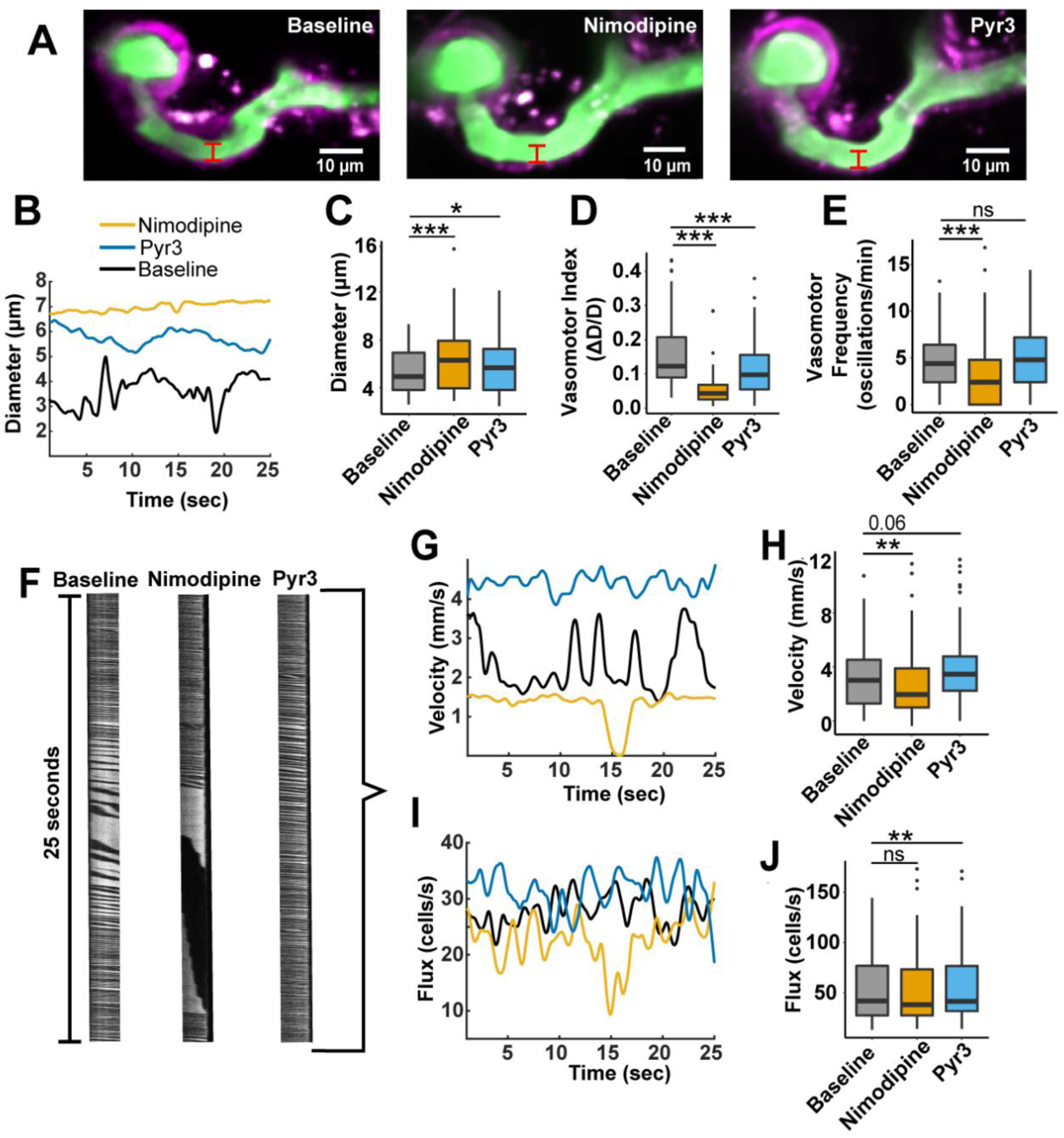
Blood vessel hemodynamics of the ACT are affected by L-type VGCC and TRPC3 channel blockers. **A)** Example of a 1st order blood vessel (green) covered by ensheathing pericytes (magenta) and the baseline diameter (red line) during nimodipine and Pyr3. **B)** Individual traces of diameter from the blood vessel in A). Diameter **(C),** vasomotor index **(D),** and vasomotion frequency **(E)** of blood vessels from the ACT covered by ensheathing pericytes. n= 101 blood vessels from N=7 mice. **F)** Representative kymographs of a 3^rd^ branch order blood vessel. The black spaces are the BCs and the white spaces are blood plasma. During nimodipine, a BC is briefly stalled, creating a long black streak. Duration of the kymographs= 30 seconds. Individual traces of BC velocity (**G)** and flux (**I**) from blood vessel kymographs of panel “F”. Velocity **(H)** and flux **(J)** of blood vessels from the ACT covered by ensheathing pericytes. n= 101 blood vessels from 7 mice. For specific p-values and mean ± SD information, please refer to SI Table S2.1 and S2.2.

A limitation when administrating nimodipine systemically is a reduction of blood pressure (35, 36). We found that nimodipine decreased blood pressure below detectable limits of a non-invasive blood pressure tail-cuff system during the two-photon imaging (Fig. S1A), likely due to hypotension-induced changes in tail volume (37). Pyr3 had no effect on blood pressure (Fig. S1A). To rule out the effect of nimodipine-induced systemic pressure changes on ensheathing pericyte Ca^2+^ and vessel hemodynamics, we performed acute *in vivo* pharmacology experiments, in which we removed the cranial window and applied nimodipine (10 μM) directly to the surface of the brain (Fig. S1B). This local drug application did not cause any difference in blood pressure compared to the sham group, and produced similar results as the systemic administration, where ensheathing pericyte Ca^2+^ signaling was reduced and vessels were dilated (Fig. S1D-F). However, BC velocity and flux increased (Fig S1G-H), similar to systemic Pyr3, which suggests that body-wide effects could cause reduced BC velocity during i.p. nimodipine administration. As such, topical application of nimodipine provides a more accurate assessment of local BC dynamics.

### L-type VGCC and TRPC3 channel blockers have different effects on specific branches of the arteriole-capillary transition zone

Recent studies have found that the kinetics of dilation/constriction differ between distinct branches of the ACT (19, 23, 24), with the first branch displaying the greatest dynamics during neurovascular coupling (23). Therefore, we evaluated the effects of systemic L-type VGCC and TRPC3 channel blockers on the 1^st^ to 3^rd^ branches of the ACT (Fig. 4A). Nimodipine dilated the 1^st^ and 2^nd^ branches but had no effect in the 3^rd^ branch (Fig. 4B). Nimodipine also decreased the vasomotor index in all three branches (Fig. 4C) and the vasomotor frequency in the 2^nd^ and 3^rd^ branches (Fig. 4D). Lastly, nimodipine tended to decrease the velocity in the 1^st^ branch (P= 0.062) but did not affect velocity in the 2^nd^ and 3^rd^ branches (Fig. 4E), nor the flux in any branch (Fig. 4F). In contrast, Pyr3 primarily affected the 2^nd^ and 3^rd^ branches, causing dilation of the 2^nd^ branch (Fig. 4B) and reduced vasomotor index in both branches (Fig. 4C). Pyr3 also increased the BC velocity and flux in the 2^nd^ and 3^rd^ branches (Fig. 4E, F), which fits with dilation of the 2^nd^ branch. Overall, these results suggest that the ACT may have a possible gradient of VGCC to TRPC3 channel activity as the branch order approaches the capillary network, which potentially influences downstream perfusion.

**Figure 4.**
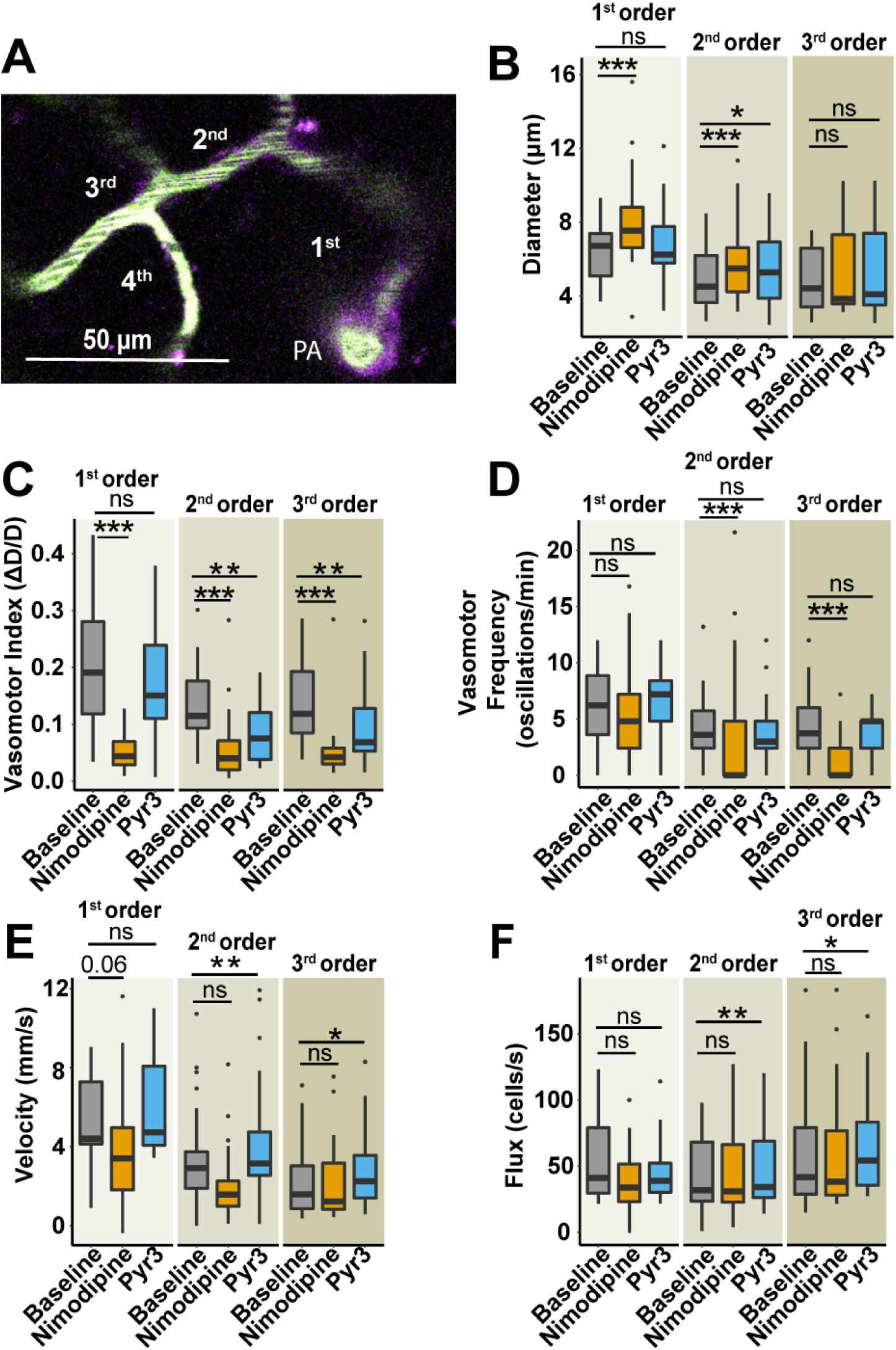
L-type VGCC and TRPC3 channel blockers affect the hemodynamics of specific branches within the arteriole transition zone. **A)** An example of the branch order classification within the ACT; ensheathing pericytes (magenta), blood vessels (green). PA= penetrating arteriole. Diameter (B), vasomotor index **(C)**, vasomotion frequency **(D)**, BC velocity **(E)** and BC flux **(F)** of 1^st^(b1), 2^nd^ (b2), and 3^rd^ (b3), branch order blood vessels. b1= 27; b2=32; b3=29 blood vessels from 7 mice. For specific p-values and mean ± SD information, please refer to SI Table S2.1 and S2.2.

### L-type VGCC and TRPC3 channel blockers affect capillary pericyte **Ca^2+^** and resting blood vessel hemodynamics in capillaries

To study the effects of VGCC and TRPC3 channels on capillary pericytes deeper in the vascular network *in vivo*, we utilized *Pdgfrb*-CreERT2:GCaMP6s^fl/fl^ mice which express inducible Cre recombinase (CreERT2) in all mural cells with platelet-derived growth factor receptor beta (*Pdgfrb;* Fig. 5A). The Cre-dependent green genetically encoded Ca^2+^ indicator, GCaMP6s, was also expressed in these cells following tamoxifen administration to activate CreERT2 (Fig. 5B, C). GCaMP6s positive capillary pericytes were identified more than 5 branches from the penetrating arteriole. Recent studies of capillary pericyte Ca^2+^ events in brain slices have found conflicting effects of L-type Ca^2+^ channel blockers and no effect of Pyr3 on Ca^2+^ event frequency (25, 26). Therefore, we conducted both brain slice pharmacology experiments with nimodipine (1 µM) and Pyr3 (3 µM; Fig. S2A-E) and *in vivo* repeated Ca^2+^ imaging like we did for ensheathing pericytes (Fig. 5B-E; same drug concentrations as Fig. 2-4). In capillary pericytes from brain slices, nimodipine had subtle effects, decreasing Ca^2+^ signal amplitude and frequency only in the processes (Fig. S2A, B). However, *in vivo* nimodipine decreased the amplitude and frequency of Ca^2+^ events in both the soma and processes of capillary pericytes (Fig. 5C-E), similar to ensheathing pericytes (Fig. 2D, E). Interestingly, Pyr3 did not affect the frequency of Ca^2+^ events in capillary pericytes in brain slices (Fig. S2C, D) or *in vivo* (Fig. 5E), similar to a recent report in brain slices (26). However, Pyr3 reduced the amplitude of Ca^2+^ events within the soma of capillary pericytes in brain slices (Fig. S2C) and in capillary pericyte processes *in vivo* (Fig. 5C, D). We also found that the area covered by individual Ca^2+^ events within capillary pericyte compartments was reduced by nimodipine and Pyr3 in both *ex vivo* and *in vivo* experiments (Fig. S2E-H), indicating that the remaining events were more localized and that these Ca^2+^ channels contribute to the spread of the Ca^2+^ events within pericytes.

**Figure 5.**
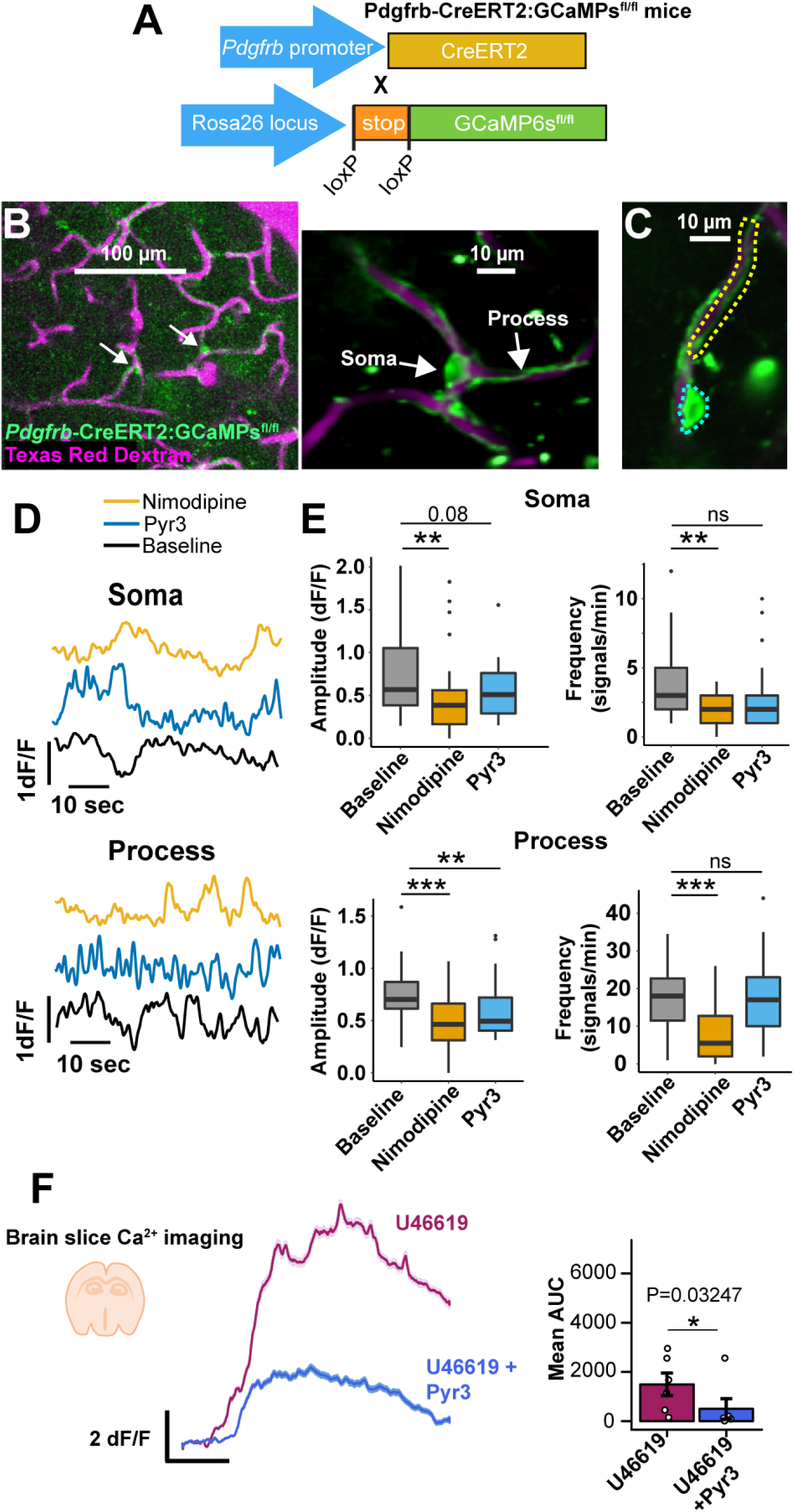
Capillary pericyte Ca^2+^ imaging during treatment with L-type VGCC and TRPC3 channel blockers. **A)** *Pdgfrb*-CreERT2:GCaMP6s^fl/fl^ mouse model. **B)** Visualization of cells expressing Cre-dependent GCaMP6s (left image, green, white arrows) and capillary pericyte morphology (right image). The vasculature was labeled using Texas red dextran (magenta). **C)** Two-photon image of a capillary pericyte morphological structures-Soma: cyan dashed line; Process; yellow dashed line. **D)** Individual Ca^2+^ signaling traces of soma and process from C). **E)** Spontaneous Ca^2+^ signaling properties (amplitude and frequency) of capillary pericyte structures, n=44 pericytes from 7 mice. **F)** Drug application to brain slices shows that Pyr3 attenuates the capillary pericyte calcium response evoked by thromboxane agonist, U46619. Left: mean ± SEM traces of the calcium response to thromboxane A2 agonist, U46619 (100nM) with and without Pyr3 (3µM). Right: The area under the curve for the calcium response from each capillary pericyte (individual dots). N= 6 mice, n= 6 pericytes. For specific p-values and mean ± SD information, please refer to SI Table S1.1 and S1.2.

TRPC3 channels are reportedly activated by Gq-GPCR signaling as a receptor-operated Ca^2+^ entry pathway (29–31). In brain slices, we applied thromboxane A2 agonist, U46619, which evoked a Ca^2+^ increase in capillary pericytes. This response to U46619 was attenuated by Pyr3 (Fig. 5F), suggesting TRPC3 channels in capillary pericytes are activated during this Gq-GPCR signaling and amplify second-messenger-induced Ca^2+^ events.

Furthermore, we evaluated capillary hemodynamics by Texas Red dextran injection (2.5%, i.v.) in the presence of the Ca^2+^ channel blockers *in vivo*, and we found that Pyr3 dilated capillaries at rest compared to baseline, while nimodipine had no dilatory effect (Fig. 6A-C). This Pyr3 dilation did not affect BC velocity or flux in capillaries compared to baseline (Fig. 6D-H). On the other hand, nimodipine decreased velocity and BC flux in capillaries (Fig. 6D-H), but also caused heterogenous fluctuations in BC velocity because of brief stalling of blood flow (Fig. 6D, E), similar to hemodynamics in the ACT (Fig. 2). These effects are potentially due to the systemic administration of nimodipine because when acutely applying nimodipine to the brain *in vivo,* there was a reduction in capillary pericyte Ca^2+^ events, without effects on capillary diameter, velocity, or flux (Fig. S3A-E). In short, these results indicate that while both TRPC3 and VGCC can contribute to capillary pericyte Ca^2+^ events, only TRPC3 channels are involved in the regulation of capillary pericyte tone.

**Figure 6:**
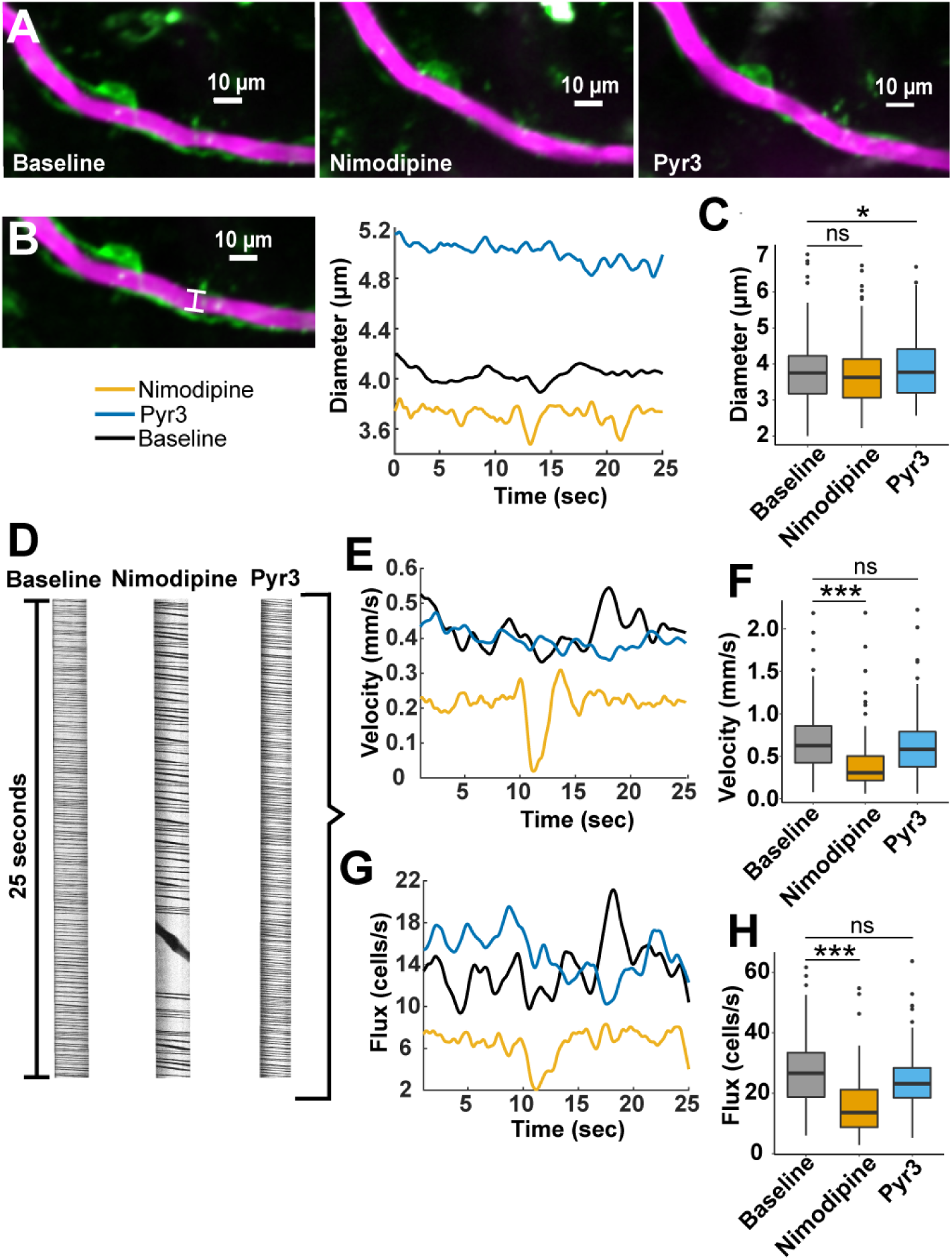
L-type VGCC and TRPC3 channel blockers have differential effects on resting capillary hemodynamics. **A)** Example of a brain capillary (magenta) covered by a capillary pericyte (green) and its fluctuation in diameter (white dashed line) in response to nimodipine and Pyr3. **B)** Two-photon image and Individual traces of brain capillary diameter from panel “A”. The continuous white line (left) represents the region where diameter was measured for the traces (right). **C)** Box plots of diameter of brain capillaries covered by capillary pericytes. **D)** Individual representative kymographs of one brain capillary. The black spaces represent BCs, and the white spaces represent blood plasma. Duration of the kymographs= 30 seconds. Individual traces of velocity **(E)** and flux (**G)** from blood vessel kymographs of panel “D”. Box plots of velocity **(F)** and flux **(H)** of brain capillaries covered by capillary pericytes. n=109 vessels from 7 mice. For specific p-values and mean ± SD information, please refer to SI Table S3.1 and S3.2.

It should be noted that we included a vehicle control (i.p. PEG400 20 mg/kg) in all of our *in vivo* experiments, which did not have effects on ensheathing or capillary pericyte Ca^2+^, vessel diameter in the ACT or capillaries, or ACT vascular dynamics (Table S12.1-S14.2). However, we found that the vehicle decreased capillary BC velocity and flux to a similar degree as nimodipine (Fig. S3F, G, Table S3.1). Since both nimodipine and Pyr3 were administered in this vehicle, it is possible that the vehicle contributed to the reduced velocity/flux observed with nimodipine and that it masked any elevations in velocity/flux after capillary dilation by Pyr3.

### L-type VGCC and TRPC3 channel blockers affect the pericyte response and hemodynamics during neurovascular coupling

To explore the influence of L-type VGCC and TRPC3 channels on pericytes during NVC, we mapped the whisker barrel cortex of mice by intrinsic optical imaging to identify regions and large pial vessels within the cranial window that responded to whisker stimulation (Fig. S4). Then, we selected ensheathing or capillary pericytes in this active area during two-photon imaging and we applied a mild electrical stimulus (500µA at 4Hz for 5s) to the whisker pad to induce neurovascular coupling. We calculated the relative change in pericyte Ca^2+^ and the percent change in blood flow during stimulation relative to the pre-stimulus baseline period (5s). In ensheathing pericytes and vessels of the ACT zone, Ca^2+^ dropped in somata and processes during the stimulation period and then slightly increased after the stimulus (overshoot; Fig. 7A; baseline session). Nimodipine and Pyr3 decreased this Ca^2+^ behaviour during the stimulus in both ensheathing pericyte compartments, but nimodipine had a stronger impact on Ca^2+^ signaling in ensheathing pericyte processes (Fig. 7A-C). In accordance with decreased Ca^2+^ in ensheathing pericytes, the electrical stimulation also dilated vessels of the ACT zone (Fig. 7D-F), with the greatest change in diameter occurring in the 1^st^ branch (Fig. 7F). We also observed a brief constriction following the stimulus (Fig. 7D), likely correlating with increased ensheathing pericyte Ca^2+^ during this period (Fig. 7A). Nimodipine and Pyr3 reduced the dilation evoked by electrical stimulation, with nimodipine having the greatest effect (Fig. 7D, E). Furthermore, nimodipine greatly reduced the dilation caused by whisker stimulation in the 1^st^ and 2^nd^ branches, but only minorly impaired the 3^rd^ branch (P=0.059; Fig. 7F). Pyr3 caused a significant reduction of the dilation caused by stimulation at the 2^nd^ branch but had no effect on the 1^st^ and 3^rd^ branches (Fig. 7F). Lastly, in control baseline conditions, BC velocity and flux increased during the period of whisker stimulation and, similar to diameter, this increase lasted longer than the stimulus period followed by an undershoot (Fig. 7G, J). Both Ca^2+^ channel blockers impaired the increase in BC velocity during stimulation, but again, nimodipine had a greater effect than Pyr3 (Fig. 7H). The effects of nimodipine and Pyr3 on velocity were obvious in the 2^nd^ and 3^rd^ branches, but not the 1st branch (Fig. 7I). Similar to BC velocity, nimodipine reduced the flux (Fig. 7J, K), particularly in the 2^nd^ and 3^rd^ branches (Fig. 7L). However, Pyr3 did not significantly reduce the flux elevation, though there was a tendency in the 2^nd^ branch for a reduction (P=0.056; Fig. 7L). Taken together, these results suggest VGCC and TRPC3 blockade attenuate the NVC response of ensheathing pericytes.

**Figure 7.**
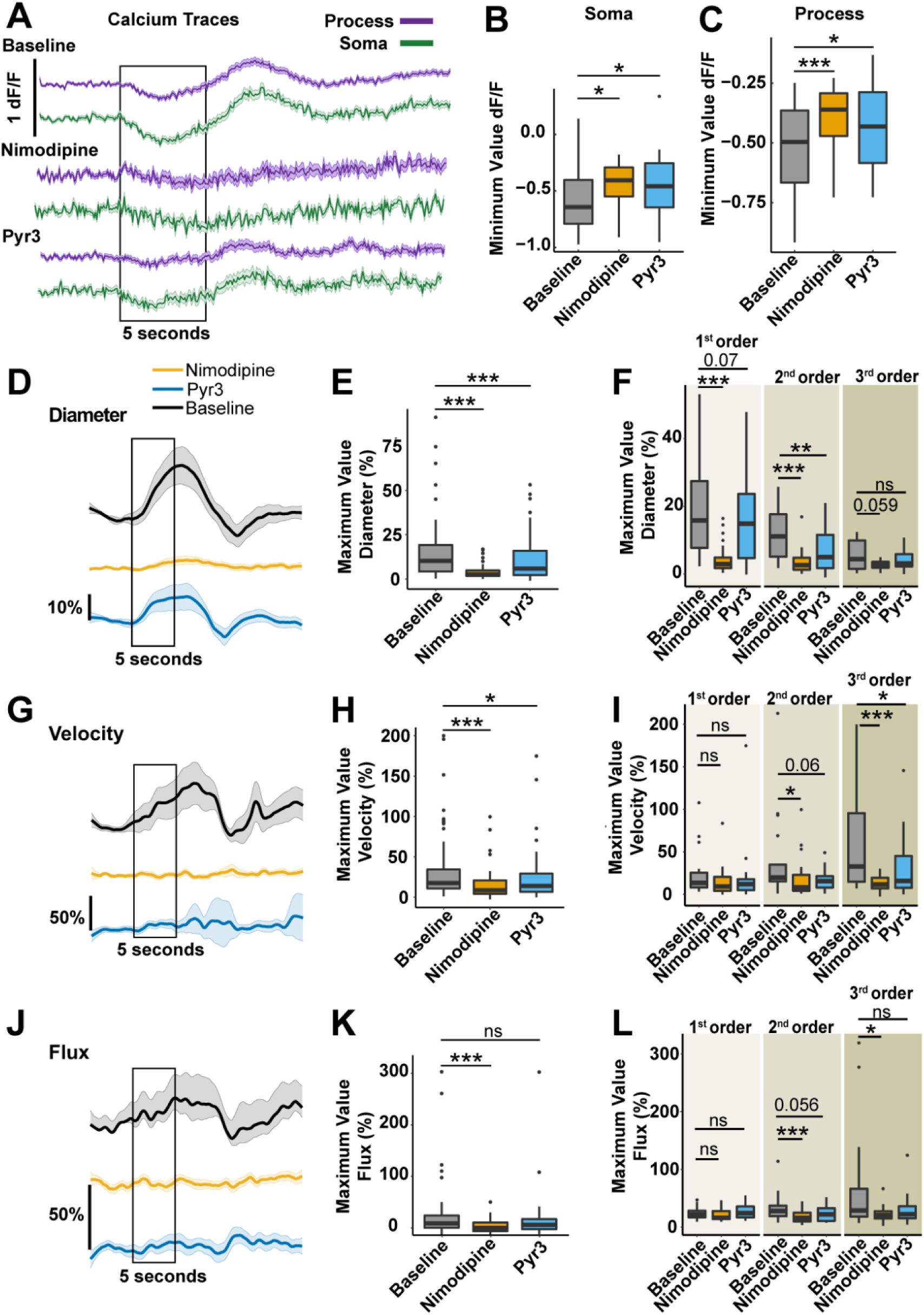
Ensheathing pericyte Ca^2+^ and hemodynamic responses during neurovascular coupling are altered by L-type VGCC and TRPC3 channel blockers. **A)** Ca^2+^ signaling traces of ensheathing pericytes during NVC. The black box represents the 5 seconds of electrical whisker stimulation. The traces show the mean value + SEM over 25s. Minimum value of Ca^2+^ drop in amplitude from ensheathing pericyte somata **(B)** and processes **(C)** during the stimulation period. n= 33 pericytes in 5 mice. Traces of the percentage change of diameter **(D)**, BC velocity **(G),** and flux **(J)** of blood vessels from the ACT during stimulation relative to the baseline period. The traces show the mean value ± SEM over 25s. n=65 vessels in 5 mice. Maximum value of diameter **(E)**, velocity **(H)** and flux **(K)** percentage change of blood vessels from the ACT during stimulation. Maximum value of diameter **(F)**, velocity **(I)** and flux **(L)** percentage change of 1^st^ (b1), 2^nd^ (b2), and 3^rd^ (b3) order blood vessels from the transition zone during stimulation. b1=16, b2=21, b3=19 vessels in 5 mice. For specific p-values and mean ± SD information, please refer to SI Table S4.1, S4.2, S5.1 and S5.2.

Capillary pericytes display a transient decrease in Ca^2+^ signaling when nearby neurons are active (6, 25). We observed a similar effect where capillary pericyte Ca^2+^ dropped during the electrical stimulation in both somata and processes, but compared to ensheathing pericytes they did not show the rise in Ca^2+^ in the post-stimulation period (Fig. 8A). Nimodipine abolished the Ca^2+^ decrease in capillary pericytes, particularly in the processes; however, Pyr3 had no effect on the response to stimulation (Fig. 8A-C). Interestingly, we detected a substantial capillary dilation during whisker pad stimulation without an undershoot of the diameter following the dilation (Fig. 8D), as clear evidence that capillaries dilate *in vivo* during neurovascular coupling. Nimodipine significantly reduced this dilation during the stimulation period, while Pyr3 had no effect (Fig. 8D, E). Likewise, electrical stimulation increased BC velocity and flux through capillaries (Fig. 8F, H). Nimodipine significantly abolished this velocity and flux behavior, whereas Pyr3 had no effect (Fig. 8F-I). This suggests that VGCC but not TRPC3 blockade diminishes the NVC response of capillary pericytes.

**Figure 8:**
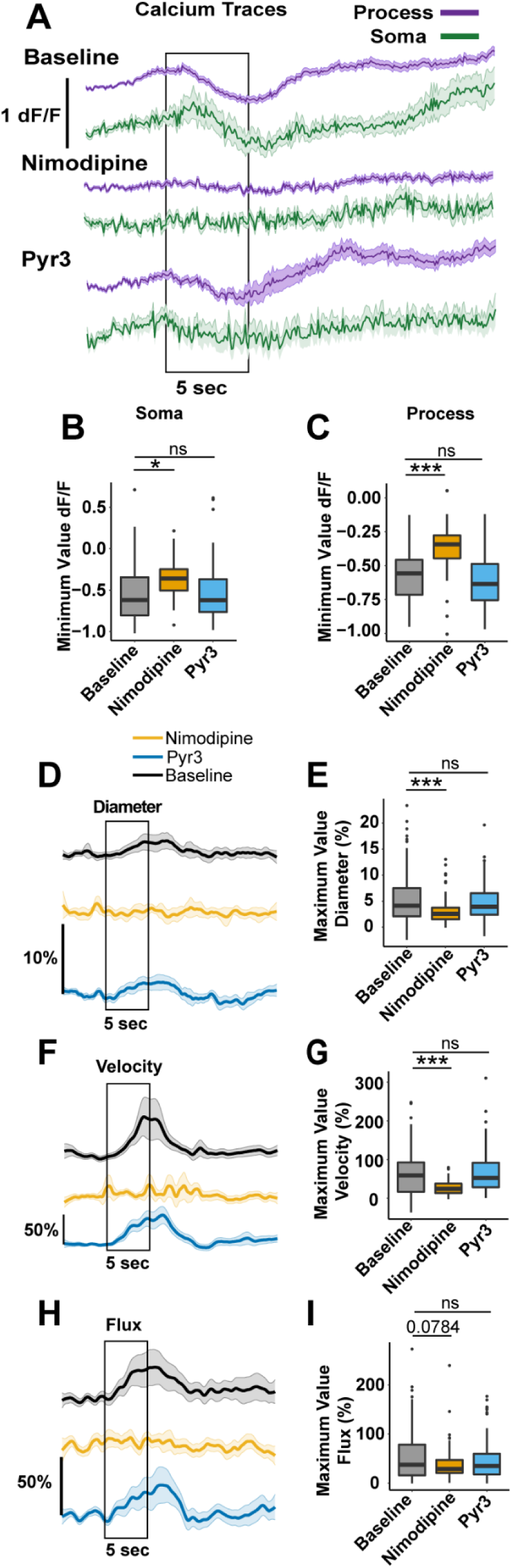
Capillary pericyte Ca^2+^ and hemodynamic responses during neurovascular coupling are altered by nimodipine, but not Pyr3. **A)** Ca^2+^ signaling traces of capillary pericytes during NVC. The black box represents the 5 seconds of electrical whisker stimulation. The traces show the mean value + SEM over the 25s. Minimum value of the Ca^2+^ drop in amplitude from capillary pericyte somata **(B)** and processes **(C),** during the stimulation period. n= 42 pericytes from 7 mice. Traces of the percentage change of diameter **(D),** BC velocity **(F)** and flux **(H)** of brain capillaries during stimulation relative to the baseline period. The traces show the mean value ± SEM over 25s. Maximum value of diameter **(E)**, velocity **(G)** and flux **(I)** percentage change in capillaries during stimulation. n=96 capillaries from 7 mice. For specific p-values and mean ± SD information, please refer to SI Table S4.1, S4.2, S6.1, S6.2.

## Discussion

Throughout this study, we identified different pericyte types by their location within the brain vasculature, but also based on their morphology and protein expression (13, 22, 24, 38). Here, we considered the impact of specific antagonists for L-type VGCC and TRPC3 channels on Ca^2+^ dynamics in different pericytes and local hemodynamics throughout the cerebrovascular tree *in vivo*. We found that 1) VGCC and TRPC3 channels are expressed by both ensheathing and capillary pericytes, 2) VGCC and TRPC3 regulate ensheathing pericyte Ca^2+^ and vasomotion events within the ACT, 3) VGCC and TRPC3 contribute to capillary pericyte Ca^2+^ events, but only TRPC3 is involved in maintaining resting capillary tone, and 4) when evoking dilation during NVC, VGCC blockade impairs the response in all vessel types, while TRPC3 blockade only affects vessels in the ACT.

Similar to recent work (25), we found that resting Ca^2+^ signaling in ensheathing pericytes occurs as synchronous waves with similar amplitude and frequency in soma and processes (Fig. 2). Nimodipine strongly reduced Ca^2+^ signaling in ensheathing pericytes (Fig. 2) and dilated vessels of the ACT, which impaired vasomotion (Fig. 3). This is comparable to smooth muscle cells of brain arterioles, where Ca^2+^ blockade by nimodipine is linked to dilation, disrupting the myosin light chain kinase (MLCK) pathway activated by the Ca^2+^-calmodulin complex (33, 39). Our results support previous reports that L-type VGCCs play a major role in ensheathing pericyte Ca^2+^ events (7, 23, 40), and contraction of vessels in the ACT (22). We also provide the first evidence that TRPC3 channels contribute to Ca^2+^ events in ensheathing pericyte processes. The effects of Pyr3 on Ca^2+^ and ACT vessel diameter were more subtle than nimodipine, but the vasomotor index was decreased (Fig. 3D) and BC flux and velocity increased (Fig. 3F-J), which suggests that TRPC3 activity contributes to ACT tone and vasomotor contractile events.

Interestingly, when evaluating the impact of these channel blockers on specific parts of the ACT, we observed unique effects on the resting diameter, vasomotion, and NVC responses of different branches (Figs. 4, 7). With nimodipine, a gradient of resting dilation occurred from the strongest response in the 1st branch to no response in the 3^rd^ branch (Fig. 4), and this was also reflected in the inhibition of NVC-vasodilation where the suppression was the strongest in the 1^st^ branch (Fig. 7). The 1^st^ branch is known to have the most robust response to vasodilatory and vasoconstrictor stimuli (23) and the fastest onset of dilation during NVC (6). Therefore, our results suggest VGCC are highly active in these mural cells, which may contribute to the vascular dynamics of this region. Reduced resting dilation in the 3^rd^ branch after nimodipine may reflect a decrease in VGCC activity (22) or a decreasing gradient of αSMA expression by pericytes (13, 23, 24, 41–43). Remarkably, nimodipine did not affect the frequency of resting vasomotor events in the 1^st^ branch, but reduced the vessel oscillations branches 2 and 3. This suggests that another mechanism drives the initiation of vasomotor events in the 1^st^ branch and VGCC are recruited to enforce stronger constriction, while in the 2^nd^ and 3^rd^ branches, VGCC activity is the principal driver of vasomotion.

When considering Pyr3, we observed resting dilation (Fig. 4) and an impairment of NVC-induced dilation (Fig. 7) in the 2^nd^ branch of the ACT, suggesting that TRPC3 regulate ensheathing pericyte tone primarily in this region. Also, Pyr3 reduced the vasomotor amplitude and elevated BC velocity and flux in the 2^nd^ and 3^rd^ branches, suggesting that TRPC3 channels are involved in contraction during vasomotion and are relevant for hemodynamics at these points in the ACT. There was no change in the frequency of vasomotor events after TRPC3 blockade, suggesting that these channels do not initiate vasomotion, but likely amplify contraction in concert with VGCC. A lack of Pyr3 effects on the 1^st^ branch of the ACT reinforces the idea that VGCC are the dominant contractile Ca^2+^ mechanism in these mural cells.

In the case of capillaries, we found that capillary pericyte Ca^2+^ signaling is more asynchronous between soma and processes (Fig. 5), similar to recent work (25). We showed for the first time *in vivo* that nimodipine decreases capillary pericyte Ca^2+^ signal amplitude and frequency in soma and processes (Fig. 5C-E; Fig. S3A, B). This contrasts *ex vivo* brain slice studies with dihydropyridines where nimodipine caused a mild Ca^2+^ frequency decrease limited to capillary pericyte somata, while nifedipine had no effect on Ca^2+^ frequency (25, 26). We also found more mild effects of nimodipine on capillary pericyte Ca^2+^ *ex vivo* particularly in processes (Fig. S2A, B), which may reflect differences in the pericyte environment in slices compared to *in vivo*. VGCC blockers are most effective when pericytes are depolarized (22), and it is possible that capillary pericyte depolarization may occur naturally *in vivo* due to physiological processes such as neurovascular unit cellular signaling and local blood flow. Nevertheless, while we observed a reduction in capillary pericyte Ca^2+^ signaling with nimodipine, there was no change in capillary diameter (Fig. 6). This suggests that VGCC do not contribute to resting capillary tone. Along the same lines, VGCC also do not play a role in pressure-induced contraction in capillaries (22).

We also found that TRPC3 activity contributes to the amplitude of capillary pericyte Ca^2+^ events *in vivo*, particularly in thin-strand processes, without changing the number of Ca^2+^ events (Fig. 5). Additionally, TRPC3 amplifies the Ca^2+^ response in brain slice capillary pericytes to Gq-GPCR agonist, U46619 (Fig. 5F). This is in-line with the idea that TRPC3 can serve as a ROC entry pathway. In other cell types, TRPC3 channels open in response to GPCR signaling and Ca^2+^ release from the endoplasmic reticulum leading to an amplification of GPCR-mediated Ca^2+^ events (44, 45). Therefore, if TRPC3 channels act via ROC in capillary pericytes, when applying Pyr3, GPCR activation would continue to trigger Ca^2+^ events and the Ca^2+^ frequency would remain the same. However, reduced Ca^2+^ influx due to TRPC3 channel blockade would decrease the amplitude of Ca^2+^ events evoked by GPCRs, which fits with our findings. Klug et al. (2023) showed that elevations in capillary pericyte Ca^2+^ in response to pressure changes are mediated by Gq-GPCR and TRPC signaling (22), and it is tempting to speculate that TRPC3 channels are involved in this pathway via ROC downstream of Gq-GPCRs.

Recent studies have suggested that pericytes can constrict capillaries and maintain capillary tone, albeit through slower mechanisms than ensheathing pericytes, possibly involving actin polymerization and Rho kinase activity (24, 43). However, specific mechanisms that contribute to this resting tone *in vivo* have not been identified. We showed that brain capillaries dilate *in vivo* after the blockade of TRPC3 channels (Fig. 6), suggesting that TRPC3-mediated mechanisms maintain capillary tone. Unexpectedly, capillary velocity and flux were unchanged by Pyr3, despite capillary dilation (Fig. 6). This could have occurred because our vehicle, PEG400, counteracted the effect of Pyr3, since we found that the vehicle decreased capillary velocity and flux compared to the baseline (Fig. S3). Despite low molecular weight PEGs like PEG400 typically being considered inert and used as drug carriers (46), the systemic application of PEG400 could potentially bind to plasma proteins due to the amphiphilic nature of PEGs (47–49), exerting more pronounced effects on brain capillaries due to their small diameter. Further research is needed to precisely understand the mechanisms underlying this observation.

During NVC, blood vessels dilate to increase flow to areas with higher energy demand owing to a number of mechanisms working in concert, such as the production of vasodilators by cells of the neurovascular unit (4) and a wave of endothelial cell hyperpolarization which passes to mural cells via gap junctions (50, 51). These mechanisms (particularly hyperpolarization) can deactivate VGCC, which causes mural cell Ca^2+^ to decrease and vessels to dilate (52). This fits with our observation that ensheathing pericyte Ca^2+^ signaling and vessel diameter within the ACT were inversely correlated, where electrical stimulation of the whisker pad caused a transient decrease in ensheathing pericyte Ca^2+^ while increasing vessel diameter and nearby BC velocity and flux (Fig. 7). Similar NVC responses were previously observed in ensheathing pericytes in the cortex and olfactory bulb (6, 9, 25). We also found a transient increase in Ca^2+^ and undershoot in vessel diameter after the stimulation period, which is known to occur in the ACT particularly within the first branch (23, 53). After nimodipine treatment, NVC responses in ensheathing pericytes were disrupted, since there was no transient drop in Ca^2+^ or change in diameter, BC velocity or flux. This suggests that the dilation during NVC cannot overcome the vessel relaxation induced by nimodipine, and blockade of VGCC may prevent VGCC deactivation in ensheathing pericytes during NVC. While nearby neurons also express VGCC, it has been shown that nimodipine (at 10 mg/kg, which is 10 times the dose used in this study) does not alter neuronal potentials evoked by whisker stimulation (54). Therefore, it seems unlikely that the reduced NVC-dilation that we observed would be due to off-target neuronal inhibition.

We also found that Pyr3 inhibition of TRPC3 activity reduced ensheathing pericyte dilation and the Ca^2+^ drop during NVC, though the effects were more subtle than nimodipine. This suggests that ensheathing pericytes still have the capacity for further dilation by NVC after the change in ensheathing pericyte tone by Pyr3 (Fig. 3). It also suggests that another mechanism other than TRPC3 inhibition, such as the deactivation of VGCC, drives the drop in Ca^2+^ and hemodynamic effects during NVC in ensheathing pericytes.

Capillary pericytes are known to decrease their Ca^2+^ signaling when nearby neurons are active (6, 25). During sensory stimulation and NVC, there is an increase in capillary diameter on a small scale (1-2%) (6, 12, 28, 55). We also found that capillaries dilated during whisker pad stimulation, but the response was larger in magnitude than previous reports (Mean ± SD; 5.677% ± 5.030%), and clearly not a “passive dilation” (6). Furthermore, we observed a considerable increase in brain capillary blood flow during NVC (Fig. 8F, H), suggesting that despite smaller-scale dilation compared to upstream vessels, there is a significant enhancement of downstream blood flow dynamics. Interestingly, VGCC blockade with nimodipine prevented the capillary pericyte Ca^2+^ decrease and dilation during NVC, which suggests that VGCC may contribute to active capillary dilation despite not impacting resting capillary pericyte tone (Fig. 6). We also found that TRPC3 blockade with Pyr3 did not impair capillary NVC, which suggests that capillaries have the capacity for further dilation despite a resting increase in diameter with Pyr3 (Fig. 6) and that other mechanisms may contribute to active capillary pericyte dilation. Collectively, these findings indicate L-type VGCCs exhibit a stronger influence in attenuating the hemodynamic responses in blood vessels within the ACT and brain capillaries, likely due to their deactivation during brain activity. In contrast, TRPC3 channels only partially affect the hemodynamic responses in blood vessels within the ACT and not the capillaries during NVC, suggesting that these channels are not primarily associated with NVC-induced hemodynamic changes.

In conclusion, the data presented in this study: 1) provide evidence of L-type VGCC and TRPC3 activity throughout pericytes of the cortical cerebrovascular network *in vivo*, 2) enhance our understanding of the role of different pericyte types in regulating local hemodynamics, and 3) provide important insight into the influence of L-type VGCC and TRPC3 channels in brain pericytes on hemodynamic responses at rest and during NVC. Our findings establish a clear distinction between brain pericyte types and their contributions to vascular tone, paving the way for novel strategies to selectively target their Ca^2+^ dynamics in controlling CBF. Future directions utilizing cell-type specific knockouts of these channels will help to fully determine their role in CBF control.

## Materials and Methods

### 1. Animals and chronic cranial window implantation

All procedures outlined below were approved by the Animal Care Committee at the University of Manitoba in accordance with the Canadian Council on Animal Care. We used two transgenic mouse lines: 1) Acta2-RCaMP1.07 mice (JAX: 028345) to visualize ensheathing pericytes since they express the Ca^2+^ indicator RCaMP1.07 in mural cells with α-SMA expression. 2) We crossed *Pdgfrb* - CreERT2 (JAX 029684 or 030201) with GCaMP6s^fl/fl^ mice (JAX 028866). Ca^2+^ indicator GCaMP6s was expressed in mural cells in these mice 2-3 weeks after tamoxifen administration (by oral gavage, 5 mg/day for 5 consecutive days). Data was collected from mice 3 to 12 months old and both sexes were used.

To access brain pericytes in mice, in an *in vivo* setting via two-photon imaging, a chronic cranial window was implanted as described in our previous work (56, 57). The surgery was done in two parts. Briefly, the mouse was anesthetized with isoflurane (4% induction, 2% maintenance) and a head post was surgically fixed at the back of the head with dental cement. In the second step 48-72 hours later, the mouse was anesthetized with a mix of medetomidine, midazolam, and fentanyl (5, 0.5, 0.05 mg/kg) and a 3x3mm piece of skull was removed and replaced with sapphire glass on the somatosensory area of the brain (Fig. 1A). The glass was permanently affixed to the skull (using dental cement). For analgesia, mice received buprenorphine slow release (0.5 mg/kg s.c.) every 72 hours and meloxicam (2 mg.kg s,c,) daily for six days following the first surgery. Mice were allowed to recover from surgery for at least three weeks before the imaging period.

### 2. Intrinsic optical imaging and selection of imaging area

Intrinsic optical imaging was done to map and select highly activated areas of the mouse somatosensory cortex with whisker stimulation. In isoflurane anesthetized animals (4% induction, 1.5% maintenance), an Ace acA2040-55um camera was used to focus and visualize the chronic cranial window area under 630nm red light produced from LEDs. Next, specific whiskers were inserted into a glass capillary connected to a piezo actuator that vibrated at a frequency of 90 Hz. During whisker stimulation, pictures of the brain surface were captured to detect the absorption of light by increased oxygenated hemoglobin in the activated area. Complete cranial window images were repeatedly captured during this process for 10-20 trials of stimulation. The images were averaged and processed to encompass the most highly activated region by selecting the darkest area (SI, Fig. S5A)

### 3. Two-photon imaging sessions

All the procedures below are described by Meza-Resillas et al. 2021 (58). Mice were anesthetized with ketamine (20 mg/kg) and xylazine (10 mg/kg) via intraperitoneal (i.p.) injection. Subsequently, 30 µL of fluorescent dextran (either fluorescein or Texas Red 70,000 MW) in saline (2.5% w/v) was injected intravenously in the tail using a catheter (Fig. 2B and 5B; (58)). Drugs were administered by i.p. injection 20 minutes prior to pericyte imaging, including nimodipine (1 mg/kg), Pyr3 (20 mg/kg) or vehicle control polyethylene glycol 400 (PEG 400). Fluorescence images were collected through the cranial window with a two-photon microscope (Bruker) with a Ti:sapphire laser set at 940 nm (GCaMP/Texas Red) or 990nm (fluorescein/RCaMP). Brain pericytes were selected based on their morphology and blood vessel branch order, saving their position as a region of interest (ROI) for future visualization and re - localization during imaging sessions with other drug conditions. Imaging parameters (i.e. laser power, zoom, PMT settings, etc.) were recorded for each ROI and applied across all imaging sessions. To capture pericyte Ca^2+^ events, movies at 128x128-pixel resolution were collected for one minute. To capture vessel hemodynamics, line scans were obtained for 30 seconds perpendicular and parallel to the vessel to generate kymographs that enabled the diameter (µm), velocity (mm/second) and BC flux (cells/second) to be calculated. We followed the same line scan drawing pattern to record hemodynamic properties in each repeated session for consistency of the measurements. To induce NVC, bipolar electrodes were percutaneously inserted into the whisker area and a STG4008 stimulus generator was used to supply electrical stimulation of 500 µA at 4 Hz for 5 seconds during the Ca^2+^ imaging movies or line scan kymographs.

### 4. Image Processing. Brain pericyte Ca^2+^ movies

Using ImageJ, an average fluorescence projection of the Ca^2+^ movie was generated to manually identify pericyte subcellular compartments (soma and process). Next, we utilized the CHIPS toolbox for MATLAB, developed by Barret et al. 2018 (59). The first step was spectral unmixing to enhance Ca^2+^ signal detection by reducing interference from the channels, such as fluorescein dextran bleeding into the red fluorescence channel (RCaMP). The Ca^2+^ analysis was run in two parts. First, Ca^2+^ traces for the regions of interest (soma/process ROIs) manually identified in ImageJ were smoothed by long-pass and band-pass filters and Ca^2+^ peaks were identified (findpeaks function) to obtain the amplitude. Second, Ca^2+^ events were automatedly detected from the movies using steps in CHIPs (based on the algorithm by Ellefsen et al. 2014 (60)) to localize pixel activity and fluorescence changes in three dimensions, using predefined thresholds to identify and group Ca^2+^ events. The total number of Ca^2+^ events that overlapped the identified soma/process ROIs was used to determine the frequency (signals/minute). **Blood vessel hemodynamic kymographs:** For diameter data collection, CHIPS algorithms were applied to the kymographs, determining diameter through full width at half maximum calculation of fluorescence. CHIPS also extracted blood vessel wall oscillation information based on diameter fluorescence fluctuations, including amplitude (vasomotor index; ΔD/D; change in diameter/mean diameter) and frequency (oscillations/minute). CHIPS radon velocity analysis algorithms were applied to calculate velocity and flux from BC activity. In some kymograph data, aberrant values were detected for single data points in the trace. Therefore, we applied thresholds based on previously reported values in literature for diameter, velocity, and flux, and replaced values that exceeded the threshold with the average of the two nearest data points. Thresholded data was interpolated and smoothed to 10 data points per second before plotting the traces.

### 5. Acute *in-vivo* pharmacology experiments

Mice with cranial windows were imaged in baseline conditions following the same procedures of section 2 and 3 to localize pericytes of interest. Next, anesthetized mice (2% isoflurane) underwent a cranial window removal surgery, in which the sapphire glass was carefully removed by drilling away the dental cement (Fig. S1B). After the window removal, a solution of 10 μM of nimodipine dissolved in PEG400 (10 µL) and cortex buffer (990 μL), or PEG400 in cortex buffer (sham solution) was topically applied with a sponge to the surface of the exposed area and left there for 20 minutes. During this time, fluorescent dextran was injected i.v. After the 20 minutes, a 2% agarose gel was applied to the brain surface and a new glass window was placed on top for two-photon imaging. Finally, ketamine (20 mg/kg) and xylazine (10 mg/kg) were administered intraperitoneally an<d isoflurane anesthesia was removed. Two-photon imaging of the mice and image processing were done using the same methodology as in sections 3 and 4. Mice from acute experiments were euthanized with a pentobarbital overdose (>150 mg/kg) after the two-photon imaging was concluded.

### 6. Blood pressure measurements

Blood pressure measurements were taken using the CODA Monitor non-invasive blood pressure system (61). We measured blood pressure of the mice during two-photon imaging sessions of repeated *in-vivo* pharmacology and acute *in-vivo* pharmacology experiments. The mice were placed on a heating pad with temperature control monitored by a rectal thermometer. The tail was inserted into the occlusion cuff and VPR systems. Blood pressure readings were taken at 15-second intervals over 5-minute periods. Each imaging session involved approximately 10 measurement periods. The measurements were manually recorded and saved for subsequent analysis using statistical software.

### 7. Brain slice Ca^2+^ imaging

Fresh mouse cortical sections from animals 3-6 months old were cut on a vibratome at 350 µm thick in ice-cold cutting buffer (N-methyl-D-glucamine, 93 mM; KCl, 3 mM; MgCl_2_ * 6H2O, 5 mM; CaCl_2_ * 2H_2_O, 0.5 mM; NaH_2_PO_4_, 1.25 mM; NaHCO_3_, 30 mM; HEPES, 20 mM; Glucose, 25 mM; Sodium Ascorbate, 5 mM; Sodium Pyruvate, 3 mM; bubbled with 95% oxygen, 5% carbon dioxide). Sections were allowed to equilibrate for 60 min at 32°C in recovery solution (NaCl, 95 mM; KCl, 3 mM; MgCl_2_ * 6H2O, 1.3 mM; CaCl_2_ * 2H2O, 2.6 mM; NaH_2_PO_4_, 1.25 mM; NaHCO_3_, 30 mM; HEPES, 20 mM; Glucose, 25 mM; Sodium Ascorbate, 5 mM; Sodium Pyruvate, 3 mM). Cortical capillary pericyte GCaMP6s fluorescence was visualized on a Ultima In Vitro Multiphoton Microscope (Bruker Fluorescence Microscopy) at 920 nm during perfusion with oxygenated aCSF (NaCl, 125 mM; KCL, 2.5 mM; NaH_2_PO_4_, 1.25 mM; MgCl_2_, 1 mM; CaCl_2_, 2 mM; NaHCO_3_, 25 mM; glucose, 25 mM) at 35°C. Ca^2+^ movies were acquired for 3 min at 256x256 pixels before and after perfusion with nimodipine (1µM) or Pyr3 (3µM). Movies were processed through MATLAB and CHIPs to identify Ca^2+^ events in different pericyte compartments, as was done for our *in vivo* analysis.

### 8. Immunostaining

Mice were injected with a pentobarbital overdose (>150 mg/kg i.p.) and monitoring closely until they reached the plane of anesthesia. They were immediately restrained on a vertical rack, their thoracic cavity exposed, and a small incision in the right artrium was made. A 21 gauge butterfly needle was inserted into the left ventricle and 20 mL of ice-cold oxygenated aCSF was perfused at a rate of ∼25 ml/min, This was followed by 60 ml of ice-cold 2% paraformaldehyde (PFA; Millipore-Sigma; P6148). Mice were then decapitated and the brains were carefully removed and fixed in ice-cold 4% PFA for 3 hours after dissection. Excess PFA was then removed from the brains and they were immediately placed in ice-cold 30% sucrose dissolved in PBS. Following 30% sucrose cryoprotection at 4°C overnight and freezing in OCT, brains were cut into 300µm thick sections on a cryostat (Leica). The sections were washed three times in TBST (0.05% Triton X-100) for 10 minutes and then permeabilized with 1% Triton in PBS (1% PBST). Then, sections were incubated with 5% Normal Donkey Serum (NDS) + 1% PBST + primary antibodies for 48 hours at 4°C. Sections were washed with 1% PBST 5 times for 30 minutes and were incubated with secondary antibodies and 1% PBST + 1% NDS for 48 hours at 4°C. Following incubation, sections were washed 3 times with 1% PBST and 2 times with PBS. Then, the slices were cleared by first dehydrating the tissue with increasing methanol concentrations: 25%, 50%, 75%, 100% for 30 min per step. Following dehydration, tissue was cleared by incubation in benzyl benzoate: benzyl alcohol (2:1) for 24 hrs. Cleared sections were placed on a coverslip and images (Z-stacks) were acquired on a confocal microscope (Zeiss LSM 810) at 63X magnification. Primary antibodies included rat-anti-CD13 (1:250, Bio-Rad; MCA2183EL), goat anti-CD31 (1:200, R&D systems, AF3628), rabbit-anti-Cav1.2 (1:200, Alomone labs, ACC-003), or rabbit anti-TRPC3 (1:200, Alomone labs, ACC-016). Secondary antibodies included donkey-anti-rat-IgG-AlexaFluor405 (1:1000, Invitrogen), donkey-anti-rabbit-IgG-AlexaFluor568 (1:1000, Invitrogen), and donkey-anti-goat-IgG-AlexaFluor647 (1:1000, Invitrogen).

### 9. Statistical Analysis

RStudio 2023.09.0+463 was used for statistics. The Normal distribution of the data was assessed visually with the Quantile-Quantile (Q-Q) plots, histograms, and box plots, and statistically with Shapiro-Wilk Test. When the data was not normally distributed, it was converted to log normal and the normal distribution was tested with the above methodology. Detection of outlier data was done using the Rosner’s test with a minimum value of 2 and a maximum value of 15 outliers for detection, the detected outliers were discarded from the data set. Linear Mix Models (LMM) were used to evaluate repeated measurements using the pharmacology condition as a fixed effect. Animal ID, pericyte ID or blood vessels ID were used as random effects. The likelihood-ratio of the LMM and their omnibus significance was performed with Anova test evaluating the LMM with and without the fixed effect. To evaluate the difference between pharmacology conditions pairwise comparisons with Holm-sequential Bonferroni correction were used. In the acute *in vivo* pharmacology study and brain slices, the distribution of results for Ca^2+^, hemodynamics, and blood pressure measurements, as well as outlier detection were assessed as mentioned above. To test the significant difference independent samples t-test was used if the data were normally distributed, and the Mann-Whitney U test as a nonparametric test when the data set did not comply with normality. Boxplots represent median and quartile values. For the brain slice Ca^2+^ analysis, plotted data represents the average of each slice. The Wilcoxon test was used to compare paired samples. Please refer to SI, Table S11 for specific information about the materials and equipment used.

## Acknowledgments

All imaging data was collected on microscopes in the Live-Cell Imaging Facility at the University of Manitoba. We would like to thank the staff in the Central Animal Care Services and Veterinary Services at the University of Manitoba for their commitment to animal care. This work was supported by Canadian Institutes for Health Research, the Manitoba Medical Service Foundation, Research Manitoba, Brain Canada through the Canada Brain Research Fund, with the financial support of Health Canada and the Azrieli Foundation. The views expressed herein do not necessarily represent the views of the Minister of Health or the Government of Canada. The Stobart lab has also received support from the Natural Sciences and Engineering Research Council of Canada (NSERC) and start-up funding from the University of Manitoba. J.M.R, M.K., and S.S. were supported by graduate studentships from the Rady Faculty of Health Sciences and Research Manitoba. S.K., J.D.R. and D.K. were supported by undergraduate research awards from the University of Manitoba, the College of Pharmacy and the College of Medicine.

