## Supplementary Figures and Tables for "L-Type Ca^2+^ channels and TRPC3 channels shape brain pericyte Ca^2+^ signaling and hemodynamics throughout the arteriole to capillary network *in vivo*"

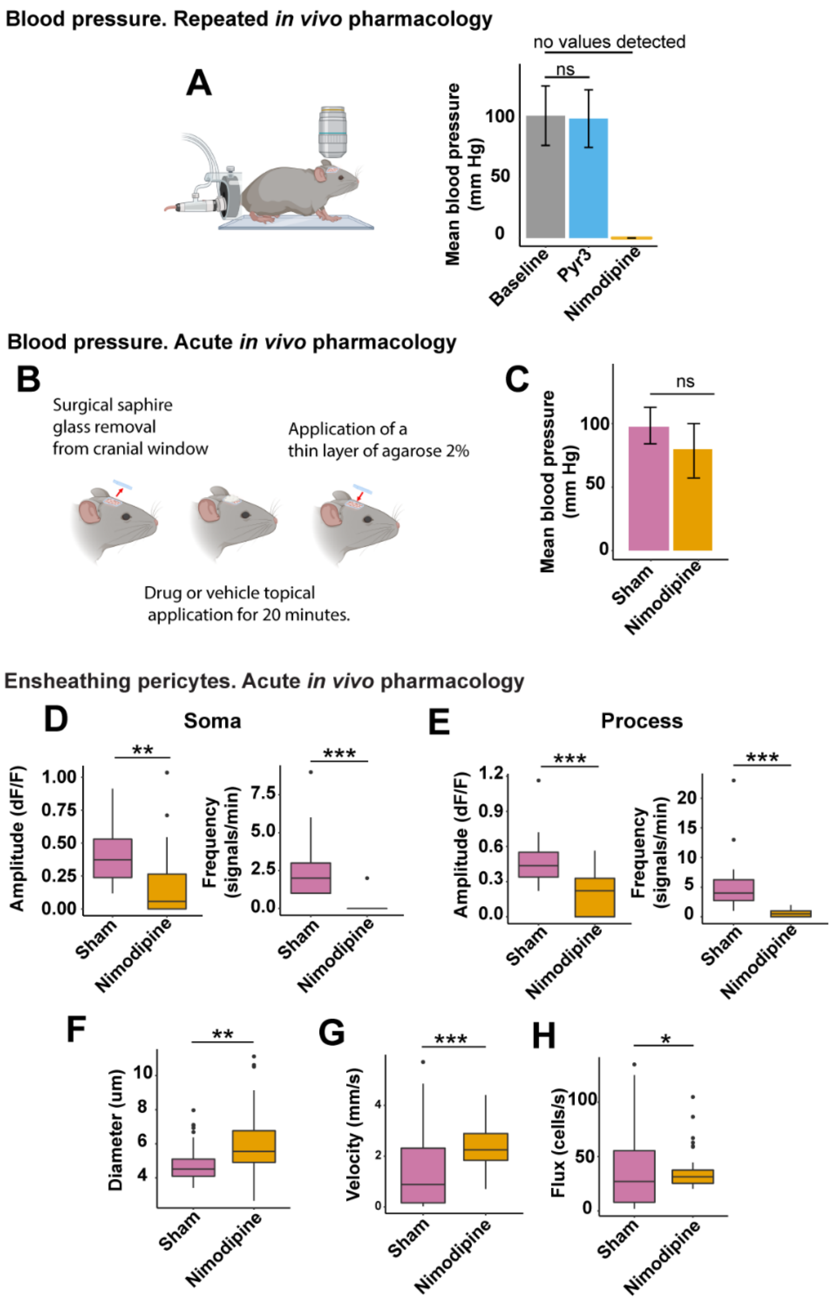

**Fig S1. Acute nimodipine application (10 µM) to the cortical surface does not affect blood pressure, but causes similar effects on ensheathing pericytes and blood flow as systemic administration. A)** Diagram of CODA monitor non-invasive blood pressure system (left). Mean blood pressure of mice measured in repeated pharmacology experiments with systemic drug administration (right). No values of blood pressure were detected with CODA system when nimodipine was applied. N= 11 mice. **B)** Acute pharmacology experiment scheme. **C)** Mean blood pressure in acute pharmacology experiments. N-Sh= 6 mice; N-nim= 9 mice. Calcium signaling properties (amplitude and frequency) of ensheathing pericyte morphology structures **D)** soma and **E)** process. NSh= 3 mice; nSh= 28 pericytes; N-nim= 4 mice; n-nim= 34 pericytes. **F)** Diameter **G)** velocity and **H)** flux of blood vessels from the transition zone covered by ensheathing pericytes.

NSh= 3 mice; nShV= 22 vessels; N-nim= 4 mice; n-nimV= 38 vessels. For specific p-values and mean ± SD information please refer to table 7-9.2.

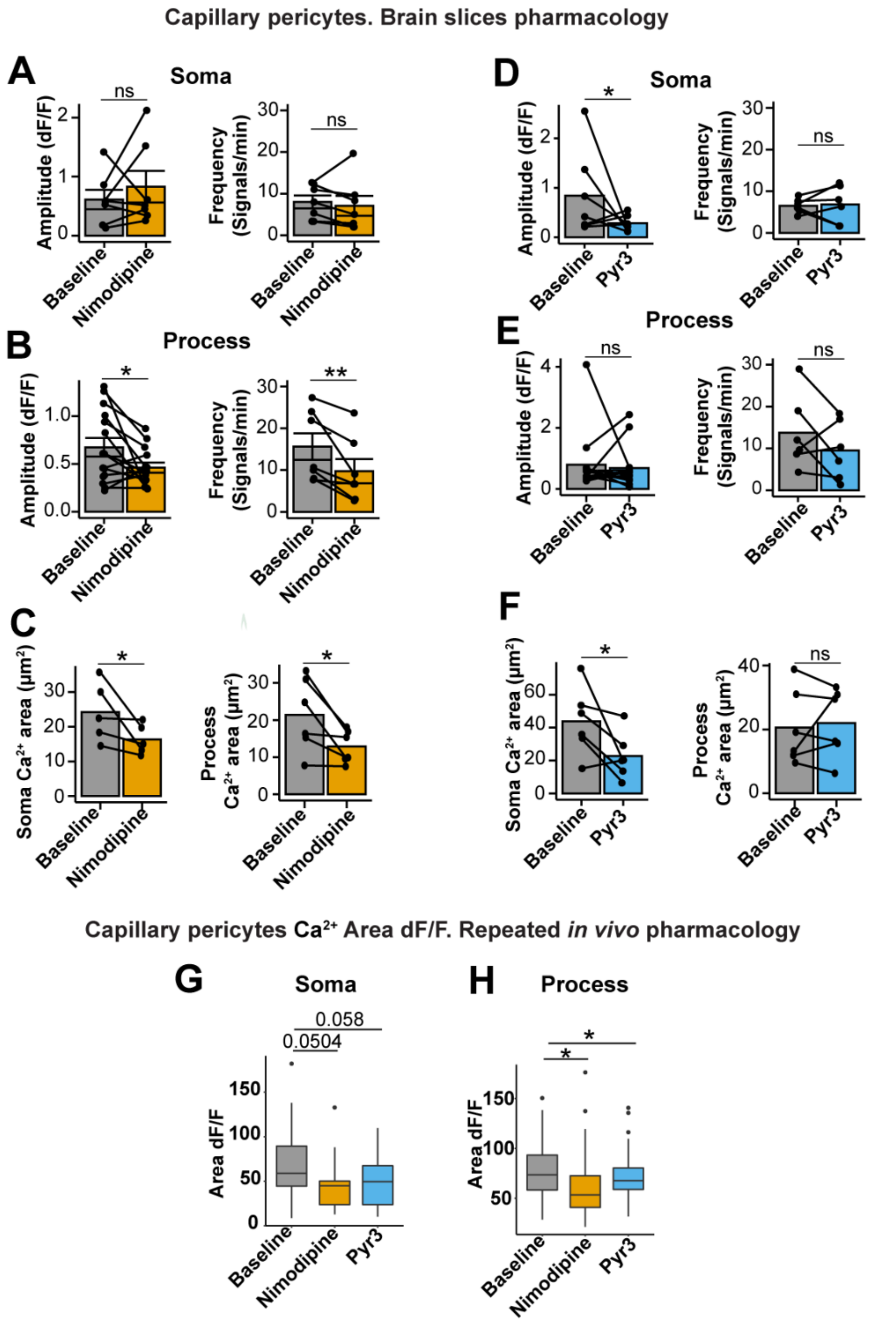

**Fig. S2. The effects of nimodipine (10µM) and Pyr3 (3µM) on capillary pericyte calcium in brain slices.** A) Nimodipine did not affect the calcium amplitude or frequency of events in capillary pericyte somata. N= 7 mice; n= 7 pericytes. B) Nimodipine reduced calcium amplitude and frequency in capillary pericyte processes. N= 7 mice; n= 14 processes from 7 pericytes. C) Nimodipine also decreased the area of calcium events within soma and processes. D) Pyr3 reduced calcium amplitude in capillary pericytes somata but did not change the frequency of calcium events. N= 6 mice; n= 6 pericytes. E) Pyr3 had no effect on calcium amplitude or frequency in processes. N= 6 mice; n= 12 processes from 6 pericytes. F) Pyr3 reduced the area of calcium events in capillary pericyte somata and processes. Box plots of calcium events area of capillary pericyte's morphology structures soma **(G)** and process **(H)** in baseline conditions and under the effect of the calcium channel blockers nimodipine, pyr3 applied systemically. N= 7; n= 44. For specific p-values and mean ± SD information please refer to tables 1.1, 1.2 and 11.1.

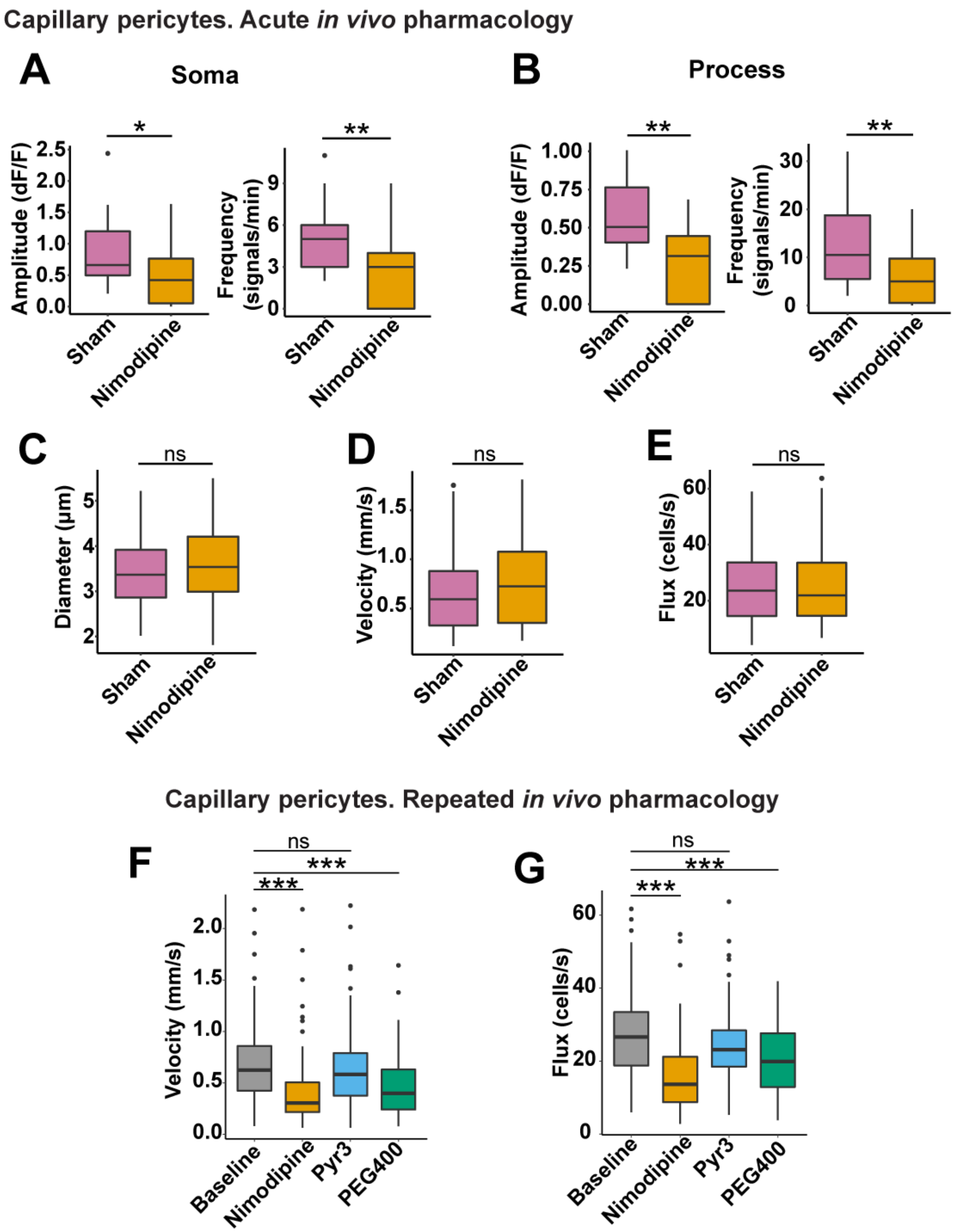

**Supporting Information. Figure S3. The effect of acute nimodipine administration or systemic vehicle control (PEG400) on capillary pericytes.** Calcium signaling properties (amplitude and frequency) of capillary pericyte morphology structures **A)** soma and **B)** process. NSh= 3 mice; nSh= 21 pericytes; N-nim= 3 mice; n-nim= 26 pericytes. **C)** Diameter **D)** velocity and **E)** flux of brain capillaries covered by capillary pericytes. NSh= 3 mice; nShV= 48 vessels; N-nim= 4 mice; n-nimV= 34 vessels. Box plots of brain capillary velocity **(F)** and flux **(G)** in baseline conditions and under the effect of the calcium channel blockers nimodipine, Pyr3 and their vehicle PEG400 applied separately via i.p. N= 7 mice; n= 109 vessels. For specific p-values and mean ± SD information please refer to tables 3.2, 8.1, 8.2, 10.1 and 10.2.

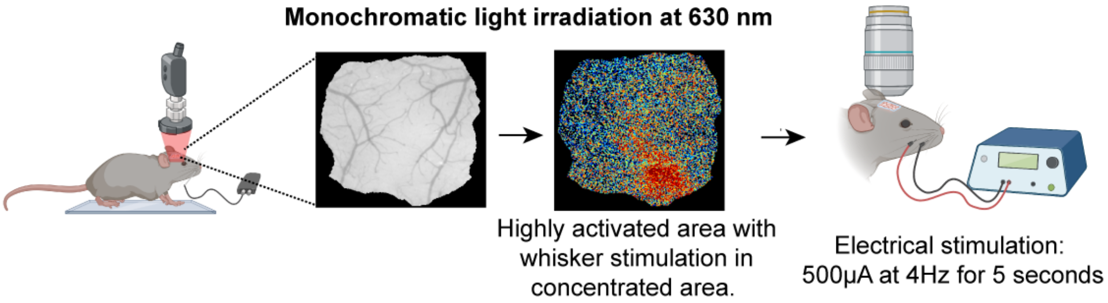

**Supporting Information. Figure S4. Intrinsic Optical imaging scheme.** Cartoon illustrating the experimental set up of intrinsic optical imaging and electrical whisker stimulation. The images of the cranial window and results from intrinsic optic imaging are an example from one mouse, 20 trials of stimulation.

**P-values and mean ± SD results of repeated *in vivo* pharmacology experiment.**

Table S1.1 P-values. Brain pericyte calcium signaling analysis. Data from Figures 2E, 5E and S2G, H.

| | Amplitude | | Frequency | | Area | |
|---|---|---|---|---|---|---|
| | Baseline-nimodipine | Baseline-Pyr3 | Baseline-nimodipine | Baseline-Pyr3 | Baseline-nimodipine | Baseline-Pyr3 |
| **Acta2-RCaMP1.07** | | | | | | |
| **Soma** | 8.40e-09*** | ns 0.319 | 1.81e-11*** | ns 0.435 | | |
| **Process** | 6.12e-08*** | ns 0.665 | 6.45e-12 *** | 0.0178* | | |
| **PDGFRβ-CreERT2:GCaMP6s$^{fl/fl}$** | | | | | | |
| **Soma** | 0.00425** | 0.0862c | 0.00472** | ns 0.180 | 0.0504c | 0.0583c |
| **Process** | 2.4e-06*** | 0.00449** | 1.96e-10*** | ns 0.947 | 0.0109* | 0.0456* |

Table S1.2. Mean ± SD. Brain pericyte calcium signaling analysis. Data from Figures 2E, 5E and S2G, H.

| | Amplitude (dF/F) | | | Frequency (signals/min) | | | Calcium event area (dF/F) | | |
|---|---|---|---|---|---|---|---|---|---|
| | Baseline | Nimodipine | Pyr3 | Baseline | Nimodipine | Pyr3 | Baseline | Nimodipine | Pyr3 |
| **Acta2-RCaMP1.07** | | | | | | | | | |
| **Soma** | 0.374± 0.207 | 0.093± 0.164 | 0.336± 0.258 | 3.964± 2.714 | 0.103± 0.409 | 4.25± 4.163 | | | |
| **Process** | 0.382± 0.260 | 0.0947 ±0.167 | 0.362± 0.282 | 6.138± 5.899 | 0.212± 0.545 | 3.611± 3.781 | | | |
| **PDGFRβ-CreERT2:GCaMP6s$^{fl/fl}$** | | | | | | | | | |
| **Soma** | 0.740± 0.429 | 0.473± 0.435 | 0.542± 0.298 | 3.581± 2.468 | 1.941± 1.344 | 2.98± 2.485 | 69.190± 34.924 | 47.448± 32.838 | 50.479± 27.602 |
| **Process** | 0.753± 0.242 | 0.505± 0.258 | 0.598± 0.258 | 17.394± 8.814 | 8± 7.635 | 17.405± 9.967 | 77.031± 28.113 | 63.113± 32.476 | 72.7±24 .987 |

Table S2.1. P-values. Blood vessels of the transition zone covered by ensheathing pericytes hemodynamic analysis. Data from Figures 3 and 4.

| | Diameter | | Vasomotor Index | | Peak frequency | |
|---|---|---|---|---|---|---|
| **Acta2-RCaMP1.07** | Baseline-nimodipine | Baseline-Pyr3 | Baseline-nimodipine | Baseline-Pyr3 | Baseline-nimodipine | Baseline-Pyr3 |
| **All blood vessels** | 9.96e-10 *** | 0.0222* | < 2e-16 *** | 2e-04 *** | 1.79e-07 *** | ns 0.797 |

| | | | | | | |
|---|---|---|---|---|---|---|
| **1st Branch** | 0.000365 *** | ns 0.361 | 5.99e-08 *** | ns 0.183 | ns 0.75 | ns 0.75 |
| **2nd Branch** | 1.89e-06 *** | 0.0199* | 2.95e-09 *** | 0.00210** | 0.000108 *** | ns 0.940 |
| **3rd Branch** | ns 0.141 | ns 0.571 | 5.59e-07 *** | 0.00861** | 0.000317 *** | ns 0.706 |
| | **Velocity** | | **Flux** | | | |
| | Baseline-nimodipine | Baseline-Pyr3 | Baseline-nimodipine | Baseline-Pyr3 | | |
| **All blood vessels** | 0.00761** | 0.0678c | ns 0.187 | 0.00458** | | |
| **1st Branch** | 0.0626c | ns 0.184 | ns 0.585 | ns 0.476 | | |
| **2nd Branch** | ns 0.113 | 0.00566** | ns 0.682 | ns 0.58 | | |
| **3rd Branch** | ns 0.192 | 0.01958* | ns 0.620 | 0.04* | | |

Table S2.2. Mean ± SD. Blood vessels of the transition zone covered by ensheathing pericytes hemodynamic analysis. Data from Figures 3 and 4.

| | **Diameter (µm)** | | | **Vasomotor Index (ΔD/D)** | | | **Peak frequency (oscillations/min)** | | |
|---|---|---|---|---|---|---|---|---|---|
| **Acta2-RCaMP1.07** | Baseline | Nimodipine | Pyr3 | Baseline | Nimodipine | Pyr3 | Baseline | Nimodipine | Pyr3 |
| **All blood vessels** | 5.247± 1.755 | 6.335± 2.591 | 5.719± 2.165 | 0.155± 0.093 | 0.055± 0.049 | 0.114± 0.081 | 4.677± 3.195 | 3.189± 4.173 | 4.784± 3.284 |
| **1st Branch** | 6.398± 1.556 | 8.061± 2.457 | 6.870± 2.025 | 0.209± 0.117 | 0.052± 0.031 | 0.172± 0.095 | 6.146± 3.524 | 5.353± 4.814 | 6.469± 3.190 |
| **2nd Branch** | 4.837± 1.689 | 5.741± 2.068 | 5.379± 1.891 | 0.136± 0.065 | 0.060± 0.060 | 0.084± 0.049 | 4.245± 2.874 | 2.488± 3.908 | 4.427± 3.686 |
| **3rd Branch** | 4.805± 1.676 | 5.337± 2.504 | 5.159± 2.338 | 0.136± 0.067 | 0.056± 0.058 | 0.100± 0.077 | 4.168± 2.864 | 1.466± 2.348 | 3.714± 2.209 |

| | **Velocity (mm/sec)** | | | **Flux (cells/sec)** | | |
|---|---|---|---|---|---|---|
| | Baseline | Nimodipine | Pyr3 | Baseline | Nimodipine | Pyr3 |
| **All blood vessels** | 3.376± 2.494 | 2.715± 2.495 | 4.069± 2.830 | 54.183± 32.012 | 55.493± 38.007 | 58.554 ±37.332 |
| **1st Branch** | 5.192± 2.361 | 3.915± 2.839 | 6.033± 2.579 | 55.739± 31.653 | 44.459± 22.826 | 47.483 ±25.687 |
| **2nd Branch** | 3.601± 2.437 | 2.278± 1.843 | 4.486± 3.213 | 53.839± 29.948 | 46.331± 30.872 | 47.795 ±31.127 |
| **3rd Branch** | 2.359± 1.897 | 2.275± 2.099 | 2.851± 1.869 | 61.367± 44.345 | 61.261± 45.978 | 66.667 ±40.130 |

Table S3.1. P-values. Blood vessels of the capillary zone covered by capillary pericytes hemodynamic analysis. Data from Figures 6C, F, H and S3 F, G.

| | Diameter | | Velocity | | Flux | |
|---|---|---|---|---|---|---|
| **PDGFRβ-CreERT2:GCaMP6s$^{fl/fl}$** | Baseline-nimodipine | Baseline-Pyr3 | Baseline-nimodipine | Baseline-Pyr3 | Baseline-nimodipine | Baseline-Pyr3 |
| **All blood vessels** | ns 0.303 | 0.0224 * | <2e-16 *** | ns 0.197 | < 2e-16 *** | ns 0.321 |
| | | | Baseline-PEG400 | | Baseline-PEG400 | |
| **All blood vessels** | | | 6.29e-13 *** | | 2.59e-06 *** | |

Table S3.2. Mean ± SD. Blood vessels of the capillary zone covered by capillary pericytes hemodynamic analysis. Data from Figures 6C, F, H and S3 F, G.

| | Diameter (µm) | | | Velocity (mm/sec) | | | Flux cells/sec) | | |
|---|---|---|---|---|---|---|---|---|---|
| **PDGFRβ-CreERT2:GCaMP6s$^{fl/fl}$** | Baseline | Nimodipine | Pyr3 | Baseline | Nimodipine | Pyr3 | Baseline | Nimodipine | Pyr3 |
| **All blood vessels** | 3.838± 0.977 | 3.808± 1.028 | 3.978± 0.986 | 0.676± 0.380 | 0.419± 0.356 | 0.677± 0.446 | 26.826± 12.032 | 16.392 ±10.014 | 24.821± 10.392 |
| | | | | PEG400 | | | PEG400 | | |
| **All blood vessels** | | | | 0.466± 0.296 | | | 20.402± 9.254 | | |

**P-values and mean ± SD of repeated *in vivo* pharmacology experiments during whisker stimulation.**

Table S4.1. P-values. Brain pericyte calcium signaling analysis during NVC. Data from Figures 7A-C and 8A-C.

| | Minimum Value | |
|---|---|---|
| | Baseline-nimodipine | Baseline-Pyr3 |
| **Acta2-RCaMP1.07** | | |
| **Soma** | 0.0179 * | 0.0347 * |
| **Process** | 0.000919 *** | 0.0108 * |
| **PDGFRβ-CreERT2:GCaMP6s$^{fl/fl}$** | | |
| **Soma** | 0.0107 * | ns0.344 |
| **Process** | 1.23e-05 *** | ns0.307 |

Table S4.2. Mean ± SD. Brain pericyte calcium signaling analysis during NVC. Data from Figures 7A-C and 8A-C.

| | **Minimum Value (dF/F)** | | |
|---|---|---|---|
| | Baseline | Nimodipine | Pyr3 |
| **Acta2-RCaMP1.07** | | | |
| **Soma** | -0.583±0.276 | -0.433±0.188 | -0.445±0.284 |
| **Process** | -0.527±0.186 | -0.40±0.119 | -0.433±0.171 |
| **PDGFRβ-CreERT2:GCaMP6s$^{fl/fl}$** | | | |
| **Soma** | -0.549±0.365 | -0.368±0.237 | -0.457±0.458 |
| **Process** | -0.570±0.205 | -0.376±0.193 | -0.608±0.198 |

Table S5.1. P-values. Blood vessels of the transition zone covered by ensheathing pericytes hemodynamic analysis during NVC. Data from Figures 7D-L.

| | **Max Value Diameter** | | **Max Value Velocity** | | **Max Value Flux** | |
|---|---|---|---|---|---|---|
| **Acta2-RCaMP1.07** | Baseline-nimodipine | Baseline-Pyr3 | Baseline-nimodipine | Baseline-Pyr3 | Baseline-nimodipine | Baseline-Pyr3 |
| **All blood vessels** | 2.35e-10 *** | 0.011 * | 0.000551*** | 0.0328 * | 5.49e-05 *** | ns 0.10805 |
| **1st Branch** | 6.53e-05 *** | ns 0.411 | ns 0.99 | ns 0.627 | ns 0.443 | ns 0.678 |
| **2nd Branch** | 5.2e-06 *** | 0.0105 * | 0.0174 * | 0.0606c | 0.000645 *** | 0.0567c |
| **3rd Branch** | 0.0592c | ns 0.258 | 0.000901 *** | 0.0305 * | 0.0175 * | ns0.1491 |

Table S5.2. Mean ± SD. Blood vessels of the transition zone covered by ensheathing pericytes hemodynamic analysis during NVC. Data from Figures 7D-L.

| | **Max Value Diameter (%)** | | | **Max Value Velocity (%)** | | | **Max Value Flux (%)** | | |
|---|---|---|---|---|---|---|---|---|---|
| **Acta2-RCaMP1.07** | Baseline | Nimodipine | Pyr3 | Baseline | Nimodipine | Pyr3 | Baseline | Nimodipine | Pyr3 |
| **All blood vessels** | 15.650± 18.032 | 3.882± 3.771 | 10.413± 12.494 | 38.144± 46.944 | 16.018 ±19.109 | 24.515± 32.379 | 41.254± 56.268 | 19.348± 12.205 | 31.018± 42.948 |
| **1st Branch** | 23.238± 20.734 | 4.693± 4.616 | 16.691± 14.760 | 25.620± 29.947 | 16.1± 20.920 | 24.096± 44.764 | 24.596± 11.906 | 21.266± 11.501 | 26.771± 12.546 |
| **2nd Branch** | 15±18.064 | 3.619± 3.678 | 9.191± 11.696 | 77.929± 101.661 | 19.87± 24.912 | 17.521± 12.614 | 33.431± 23.891 | 17.610± 9.814 | 23.568± 13.212 |
| **3rd Branch** | 5.492± 4.4790 | 2.637± 1.426 | 3.766± 2.888 | 80.616± 82.611 | 13.351 ±9.599 | 33.078± 36.763 | 71.567± 93.544 | 22.355± 15.146 | 29.747± 26.822 |

Table S6.1. P-values. Blood vessels of the capillary zone covered by capillary pericytes hemodynamic analysis during NVC. Data from Figures 8D-I.

| | **Max Value Diameter** | | **Max Value Velocity** | | **Max Value Flux** | |
|---|---|---|---|---|---|---|
| **PDGFRβ-CreERT2:GCaMP6s$^{fl/fl}$** | Baseline-nimodipine | Baseline-Pyr3 | Baseline-nimodipine | Baseline-Pyr3 | Baseline-nimodipine | Baseline-Pyr3 |

| | | | | | | |
|---|---|---|---|---|---|---|
| **All blood vessels** | 0.00248 ** | ns0.330 83 | 3.85e-09 *** | ns0.819 | ns 0.102 | ns0.909 |

Table S6.2 Mean ± SD. Blood vessels of the capillary zone covered by capillary pericytes hemodynamic analysis during NVC. Data from Figures 8D-I.

| | **Max Value Diameter (%)** | | | **Max Value Velocity (%)** | | | **Max Value Flux (%)** | | |
|---|---|---|---|---|---|---|---|---|---|
| **PDGFRβ-CreERT2:GCaMP6s$^{fl/fl}$** | Baseline | Nimodipine | Pyr3 | Baseline | Nimodipine | Pyr3 | Baseline | Nimodipine | Pyr3 |
| **All blood vessels** | 5.677± 5.030 | 3.368± 2.774 | 4.727± 3.594 | 67.729± 57.556 | 27.378± 18.563 | 69.306± 58.596 | 52.477± 49.349 | 37.953± 33.898 | 49.551± 45.553 |

**P-values and mean ± SD Results of blood pressure measurements.**

Table S7. P-values of the mean blood pressure in repeated and acute pharmacology experiments. Data from Figures S1A, C.

| | **P-Value** | **P-Value** |
|---|---|---|
| **Repeated pharmacology experiments** | Baseline-nimodipine | Baseline-pyr3 |
| | NO VALUES DETECTED | ns 0.751 |
| | **P-Value** | |
| **Acute pharmacology experiments** | Sham-Nimodipine | |
| | ns0.103 | |

**P-values and mean ± SD Results of acute *in vivo* pharmacology experiments.**

Table S8.1. P-values. Brain pericyte calcium signaling analysis. Data from Figures S1D, E and S4A, B.

| | **Amplitude** | **Frequency** |
|---|---|---|
| | Sham-nimodipine | Sham-nimodipine |
| **Acta2-RCaMP1.07** | | |
| **Soma** | 0.00262** | 0.000367 *** |
| **Process** | 0.0002 *** | 2.831e-05 *** |

| PDGFRβ-CreERT2:GCaMP6s$^{fl/fl}$ | | |
|---|---|---|
| **Soma** | 0.01181* | 0.00166 ** |
| **Process** | 0.00158** | 0.00939 ** |

Table S8.2. Mean ± SD. Brain pericyte calcium signaling analysis. Data from Figures S1D, E and S4A, B.

| | **Amplitude (dF/F)** | | **Frequency (signals/min)** | |
|---|---|---|---|---|
| | Sham | Nimodipine | Sham | Nimodipine |
| **Acta2-RCaMP1.07** | | | | |
| **Soma** | 0.390±0.215 | 0.195±0.283 | 2.8±2.210 | 0.444±0.881 |
| **Process** | 0.471±0.209 | 0.190±0.193 | 5.687±5.582 | 0.6±0.699 |
| **PDGFRβ-CreERT2:GCaMP6s$^{fl/fl}$** | | | | |
| **Soma** | 0.936±0.659 | 0.472±0.439 | 5±2.380 | 2.666±2.566 |
| **Process** | 0.573±0.242 | 0.296±0.226 | 12.444±8.276 | 5.954±5.802 |

Table S9.1. P-values. Blood vessels of the transition zone covered by ensheathing pericytes hemodynamic analysis. Data from Figures S1F-H.

| | **Diameter** | **Velocity** | **Flux** |
|---|---|---|---|
| **Acta2-RCaMP1.07** | Sham-nimodipine | Sham-nimodipine | Sham-nimodipine |
| **All blood vessels** | 0.00295 ** | 8.794e-06*** | 0.0216* |

Table S9.2. Mean ± SD. Blood vessels of the transition zone covered by ensheathing pericytes hemodynamic analysis. Data from figure S1F-H.

| | **Diameter (µm)** | | **Velocity (mm/sec** | | **Flux (cells/sec)** | |
|---|---|---|---|---|---|---|
| **Acta2-RCaMP1.07** | Sham | Nimodipine | Sham | Nimodipine | Sham | Nimodipine |
| **All blood vessels** | 4.789±1.045 | 6.017±2.187 | 1.617±1.976 | 2.572±1.219 | 34.893±33.488 | 37.319±18.651 |

Table S10.1. P-values. Blood vessels of the capillary zone covered by capillary pericytes hemodynamic analysis. Data from Figures S3C-E.

| | **Diameter** | **Velocity** | **Flux** |
|---|---|---|---|
| **PDGFRβ-CreERT2:GCaMP6s$^{fl/fl}$** | Sham-nimodipine | Sham-nimodipine | Sham-nimodipine |
| **All blood vessels** | ns 0.4091 | ns 0.4617 | ns 0.8023 |

Table S10.2. Mean ± SD. Blood vessels of the capillary zone covered by capillary pericytes hemodynamic analysis. Data from Figures S3C-E.

| | **Diameter (µm)** | **Velocity (mm/sec)** | **Flux cells/sec)** |
|---|---|---|---|

| PDGFRβ-CreERT2:GCaMP6s$^{fl/fl}$ | Sham | Nimodipine | Sham | Nimodipine | Sham | Nimodipine |
|---|---|---|---|---|---|---|
| **All blood vessels** | 3.431±0.740 | 3.579±0.833 | 0.694±0.421 | 0.773±0.455 | 26.126±15.190 | 26.751±15.640 |

**P-values and mean ± SD Results of brain slices (*ex-vivo* pharmacology) experiments.** Table S11.1. P-values. Brain slices of capillary pericytes calcium signaling analysis. Data from Figures S2A-F.

| | **Amplitude** | | **Frequency** | | **Area** | |
|---|---|---|---|---|---|---|
| | Baseline-nimodipine | Baseline-Pyr3 | Baseline-nimodipine | Baseline-Pyr3 | Baseline-nimodipine | Baseline-Pyr3 |
| **PDGFRβ-CreERT2:GCaMP6s$^{fl/fl}$** | | | | | | |
| **Soma** | ns 0.267 | 0.04* | ns 0.1484 | ns 0.656 | 0.03125* | 0.03125* |
| **Process** | 0.0488* | ns 0.469 | 0.0078** | ns 0.1563 | 0.0156* | ns0.5 |

**P-values and mean ± SD results of repeated *in vivo* pharmacology experiment comparing baseline and vehicle PEG400 conditions.**

Table S12.1 P-values. Brain pericyte calcium signaling analysis with PEG400.

| | **Amplitude** | **Frequency** | **Area** |
|---|---|---|---|
| | Baseline-PEG400 | Baseline-PEG400 | Baseline-PEG400 |
| **Acta2-RCaMP1.07** | | | |
| **Soma** | ns 0.363 | ns 1.000 | |
| **Process** | ns 1.000 | ns 0.135 | |
| **PDGFRβ-CreERT2:GCaMP6s$^{fl/fl}$** | | | |
| **Soma** | ns 0.477 | ns 0.769 | ns 0.415 |
| **Process** | ns 0.236 | ns 1.000 | ns 1.000 |

Table S12.2. Mean ± SD. Brain pericyte calcium signaling analysis with PEG400.

| | **Amplitude (dF/F)** | | **Frequency (signals/min)** | | **Area (dF/F)** | |
|---|---|---|---|---|---|---|
| | Baseline | PEG400 | Baseline | PEG400 | Baseline | PEG400 |
| **Acta2-RCaMP1.07** | | | | | | |
| **Soma** | 0.374± 0.207 | 0.276± 0.207 | 3.964± 2.714 | 4.647± 4.591 | | |

| | | | | | | |
|---|---|---|---|---|---|---|
| **Process** | 0.382± 0.260 | 0.326± 0.246 | 6.138± 5.899 | 3.904± 4.134 | | |
| **PDGFRβ-CreERT2:GCaMP6s$^{fl/fl}$** | | | | | | |
| **Soma** | 0.740± 0.429 | 0.604± 0.354 | 3.581± 2.468 | 3.283± 2.273 | 69.190± 34.924 | 57.595± 39.206 |
| **Process** | 0.753± 0.242 | 0.696± 0.285 | 17.394± 8.814 | 17.884± 9.672 | 77.031± 28.113 | 76.559± 27.375 |

Table S13.1 P-values. Blood vessels of the transition zone covered by ensheathing pericytes hemodynamic analysis with PEG400.

| | **Diameter** | **Vasomotor Index** | **Peak Frequency** |
|---|---|---|---|
| **Acta2-RCaMP1.07** | Baseline-PEG400 | Baseline-PEG400 | Baseline-PEG400 |
| **All blood vessels** | ns 0.979 | ns 0.117 | ns 0.861 |
| **1st Branch** | ns 0.517 | ns 0.845 | ns 1.000 |
| **2nd Branch** | ns 0.153 | ns 0.306 | ns 1.000 |
| **3rd Branch** | ns 1.000 | ns 0.299 | ns 1.000 |

| | **Velocity** | **Flux** |
|---|---|---|
| | Baseline-PEG400 | Baseline-PEG400 |
| **All blood vessels** | ns 0.642 | ns 0.815 |
| **1st Branch** | ns 0.561 | ns 0.696 |
| **2nd Branch** | ns 0.398 | ns 1.000 |
| **3rd Branch** | ns 0.487 | ns 1.000 |

Table S13.2 Mean ± SD. Blood vessels of the transition zone covered by ensheathing pericytes hemodynamic analysis with PEG400.

| | **Diameter (μm)** | | **Vasomotor Index (ΔD/D)** | | **Peak Frequency (oscillations/min)** | |
|---|---|---|---|---|---|---|
| **Acta2-RCaMP1.07** | Baseline | PEG400 | Baseline | PEG400 | Baseline | PEG400 |
| **All blood vessels** | 5.247± 1.755 | 5.186± 1.946 | 0.155 ± 0.0937 | 0.131 ± 0.0845 | 4.677 ± 3.195 | 5.500 ± 4.482 |

| | | | | | | |
|---|---|---|---|---|---|---|
| **1st Branch** | 6.398 ± 1.556 | 5.966± 1.956 | 0.209± 0.117 | 0.199± 0.119 | 6.146 ± 3.524 | 7.200 ± 5.299 |
| **2nd Branch** | 4.837± 1.689 | 4.983± 1.687 | 0.136± 0.065 | 0.114± 0.067 | 4.245± 2.874 | 4.235± 3.652 |
| **3rd Branch** | 4.805± 1.676 | 4.660± 2.236 | 0.136± 0.067 | 0.104± 0.085 | 4.168± 2.864 | 3.490± 2.707 |

| | **Velocity (mm/sec)** | | **Flux (cells/sec)** | |
|---|---|---|---|---|
| | Baseline | PEG400 | Baseline | PEG400 |
| **All blood vessels** | 3.376± 2.494 | 3.233± 1.920 | 54.183± 32.012 | 42.314± 18.718 |
| **1st Branch** | 5.192± 2.361 | 4.011± 1.550 | 55.739± 31.653 | 40.289± 18.908 |
| **2nd Branch** | 3.601± 2.437 | 3.608± 2.051 | 53.839± 29.948 | 42.415± 21.185 |
| **3rd Branch** | 2.359± 1.897 | 2.034± 1.628 | 61.367± 44.345 | 41.954± 16.302 |

Table S14.1 P-values. Blood vessels of the capillary zone covered by capillary pericytes hemodynamic analysis with PEG400.

| | **Diameter** | **Velocity** | **Flux** |
|---|---|---|---|
| **PDGFRβ-CreERT2:GCaMP6s$^{fl/fl}$** | Baseline-PEG400 | Baseline-PEG400 | Baseline-PEG400 |
| **All blood vessels** | ns 0.861 | 6.29e-13 *** | 2.59e-06 *** |

Table S14.2 Mean ± SD. Blood vessels of the capillary zone covered by capillary pericytes hemodynamic analysis with PEG400.

| | **Diameter (µm)** | | **Velocity (mm/sec)** | | **Flux (cells/sec)** | |
|---|---|---|---|---|---|---|
| **PDGFRβ-CreERT2:GCaMP6s$^{fl/fl}$** | Baseline | PEG400 | Baseline | PEG400 | Baseline | PEG400 |
| **All blood vessels** | 3.838± 0.977 | 3.825± 0.936 | 0.676± 0.380 | 0.466± 0.296 | 26.826± 12.032 | 20.402± 9.254 |

ns = no significant difference
P-value<0.05=*; P-value<0.01=**; P-value<0.001=***.
A letter "c" besides the numerical value indicates that the p-value is close to a significant difference.
NA= No applicable

**Materials and Equipment used.**

Table 11. Materials and Equipment used.

| Material/ Equipment | Company | Catalog Number | Comments |
|---|---|---|---|
| Acta2-RCaMP1.07 mice | The Jackson Laboratory | 28345 | Common Name: CHROMus line acta2-RCaMP1.07 |
| Applicators (Regular) | Bisco | X-80250P | |
| Agarose | Sigma Aldrich | | |
| ace acA2040-55um camera | Basler | 107210 | |
| BioFormats package for MATLAB | NA | NA | Available in: https://docs.openmicroscopy.org/bioformats/ |
| CHIPS MATLAB toolbox | NA | NA | Barrett MJP, Ferrari KD, Stobart JL, Holub M, Weber B. CHIPS: an Extensible Toolbox for Cellular and Hemodynamic Two-Photon Image Analysis. Neuroinformatics. 2018;16(1):145-147. doi:10.1007/s12021-017-9344-y. Available in: https://github.com/EIN-lab/CHIPS |
| Clear Ultrasound Gel, Medium viscosity | HealthCare Plus | UGC250 | |
| Dextran, fluorescein, 70,000 MW, anionic | Thermo Fisher Scientific | D1823 | |
| Dextran, Texas Red, 70,000 MW, neutral | Thermo Fisher Scientific | D1830 | |
| Dental cement | Bisco dental | NA | |
| Eye Lube Plus | Optixcare | NA | |
| FIJI | Image J | NA | Available in: https://imagej.net/Fiji/Downloads |
| GCaMP6s$^{fl/fl}$ mice | The Jackson Laboratory | 28866 | Common Name: Ai96(RCL-GCaMP6s) (C57BL/6J) or Ai96 (C57BL/6J) |
| Head Post | NA | NA | This product is custom made |
| Head Post fixing platform | University of Zurich | NA | |
| Isoflurane | University of Manitoba | NA | |

| | | | |
|---|---|---|---|
| Ketamine (Narketan 100 mg/mL) | Vetoquinol | 440893 | |
| MATLAB R2020b | Mathworks | NA | Available in: https://www.mathworks.com/downloads/ |
| MC_Stimulus II | Multichannel systems | NA | Please refer: https://www.multichannelsystems.com/sites/multichannelsystems.com/files/documents/manuals/MCS_STG4004%2BSTG4008_Manual.pdf |
| 30 G Needle 0.3mmx25mm | BD PrecisionGlide | 305128 | |
| Nimodipine | Sigma Aldrich | 66085-59-4 | |
| Objective XLUMPLFLN20XW | Olympus | NA | Please refer: https://www.olympus-lifescience.com/en/objectives/lumplfln-w/ |
| PDGFRβ-CreERT2 mice | The Jackson Laboratory | 30201 | Common Name: PDGFRβ-P2A-CreER$^{T2}$ |
| Polyethylene Tubing, PE10 I.D. 28mm (0.11”) O.D. 61mm (.024”) | BD Intramedic | 427401 | |
| Pyr3 | TOCRIS | 3751 | |
| Prairie View | Bruker Fluorescence Microscopy | NA | Please refer: https://www.bruker.com/en/products-and-solutions/fluorescence-microscopy/multiphoton-microscopes/ultima-in-vitro.html |
| Polychrome IV | Till photonics | NA | |
| RStudio 2023.09.0+463 | Posit | NA | Available in: https://posit.co/download/rstudio-desktop/ |
| Saphire glass | NA | NA | This product is custom made |
| STG4008 stimulus generator | Multichannel systems | NA | Please refer: https://www.multichannelsystems.com/sites/multichannelsystems.com/files/documents/manuals/MCS_STG4004%2BSTG4008_Manual.pdf |
| Tamoxifen | Sigma Aldrich | 10540-29-1 | |
| Ultima In Vitro Multiphoton Microscope | Bruker Fluorescence Microscopy | NA | Please refer: https://www.bruker.com/en/products-and-solutions/fluorescence-microscopy/multiphoton-microscopes/ultima-in-vitro.html |
| Under Tank Heater | Reptitherm U.T.H | E169064 | |

| | | |
|---|---|---|
| Xylazine (Rompun 20 mg/mL) | Bayer HealthCare | 2169592 |
